# Programmed mechano-chemical coupling in reaction-diffusion active matter

**DOI:** 10.1101/2021.03.13.435232

**Authors:** Anis Senoussi, Jean-Christophe Galas, André Estevez-Torres

**Affiliations:** Sorbonne Université, CNRS, Institut de Biologie Paris-Seine (IBPS), Laboratoire Jean Perrin (LJP), F-75005, Paris

**Keywords:** DNA programming, active matter, morphogenesis, pattern formation, life-like material

## Abstract

Embryo morphogenesis involves a complex combination of pattern-forming mechanisms. However, classical *in vitro* patterning experiments explore only one mechanism at a time, thus missing coupling effects. Here, we conjugate two major pattern-forming mechanisms —reaction-diffusion and active matter— by integrating dissipative DNA/enzyme reaction networks within an active gel composed of cytoskeletal motors and filaments. We show that the strength of the flow generated by the active gel controls the mechano-chemical coupling between the two subsystems. This property was used to engineer a synthetic material where contractions trigger chemical reaction networks both in time and space, thus mimicking key aspects of the polarization mechanism observed in *C. elegans* oocytes. We anticipate that reaction-diffusion active matter will promote the investigation of mechano-chemical transduction and the design of new materials with life-like properties.

---

Living embryos get their shape through a complex combination of chemical and physical processes that take place out of equilibrium. Basically, biochemical reaction networks process information at particular points of space and time, while active gels generate mechanical forces and flows.^1–3^ These two generic processes are intertwined through diverse chemo-mechanical and mechano-chemical couplings and both continuously consume chemical energy. For instance, the pathway Rho GTPase brings about local contractions of the actomyosin cortex in *Xenopus*,^4^ which regulates the cell cycle, while cytoplasmic flows of this same cortex trigger the PAR reaction network in *C. elegans*, inducing embryo polarization.^5^

The development of *in vitro* dissipative molecular systems is key to produce non-equilibrium materials and to investigate these processes in a controlled environment for testing theoretical predictions.^6,7^ On the chemical side, *in vitro* out-of-equilibrium reaction networks produce spatio-temporal concentration patterns through reaction or reaction-diffusion instabilities.^8–15^ On the mechanical side, active gels made of cytoskeletal motors and filaments reconstituted *in vitro* ^6,16–25^ convert chemical energy into mechanical work and generate static patterns and flows through hydrodynamic instabilities.^26^ More recently, efforts have focused on coupling reaction networks with active fluids and gels, resulting in redox-controlled self-oscillating gels^27^ and DNA-controlled passive^28^ and active gels.^29–31^ However, so far, only one of the two systems could be maintained out of equilibrium, either the reaction network, ^27^ or the gel,^28–32^ and thus only one of them could exhibit spatio-temporal self-organization. This constraint suppresses the rich variety of mechano-chemical couplings that are essential for pattern generation during development.^1–3^ It thus constitutes a major limitation for the design of self-shaping materials inspired by embryogenesis.^7,33,34^

To create a functional mechano-chemical dissipative material we assembled two dissipative molecular subsystems, a chemical and a mechanical one. The chemical subsystem is made of a network of DNA/enzyme reactions that produce single-stranded DNA (ssDNA) molecules^14^ (Figure 1a). The mechanical subsystem is an active gel composed of bundles of protein filaments propulsed by molecular motors^16,18^ (Figure 1b). Each subsystem is maintained out of equilibrium *via* the hydrolysis of high-energy compounds, respectively deoxynucleosidetriphosphates (dNTPs) and adenosinetriphosphate (ATP).

**Figure 1.**
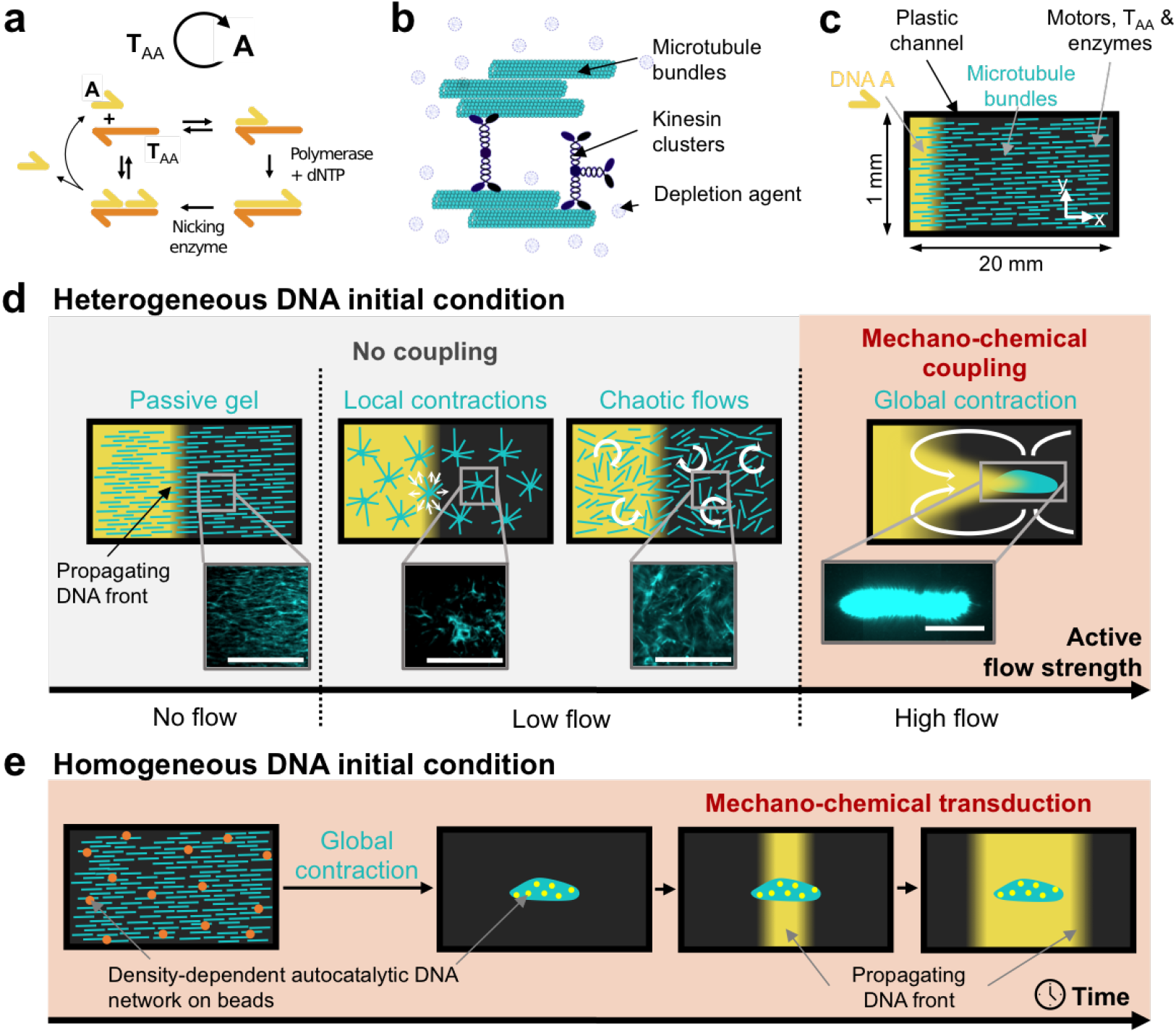
In a reaction-diffusion active matter system, the mechano-chemical coupling is controlled by the strength of the active flow and it can be used to engineer mechano-chemical transduction in a synthetic material. (a) Scheme of the chemical subsystem involving the autocatalytic amplification of DNA strand **A** in the presence of enzymes and template strand **T**_AA_. Harpoon-ended arrows denote ssDNA. (b) Cartoon of the mechanical subsystem: an active gel formed by microtubules bundled together by a depletion agent and clusters of kinesin-1 motors. (c) Sketch of the channel in which the DNA front propagation and the active gel dynamics were observed by fluorescence microscopy. In most experiments, **A** (yellow) is initially present only on the left side and the microtubule bundles (light blue) are aligned along *x*. (d) A mechano-chemical coupling between the two subsystems is achieved by increasing the strength of the flows generated by the active gel, which induces four different microtubule structures (light blue) and two DNA patterns (yellow). The white arrows represent the hydrodynamic flows generated by the active gel. Fluorescence images of the microtubules are represented for each morphology. Scale bars are 0.5 mm. (e) When the initial condition of DNA **A** is homogeneously distributed and the kinetics of DNA auto-catalysis depend on the density of the active gel, gel contraction is transduced into a DNA propagating front starting at the position of the contracted gel.

The chosen chemical subsystem has four advantages. Firstly, due to DNA hybridization rules, it can be easily reprogrammed into a variety of dissipative dynamics such as oscillations,^35^ bistability and excitability. ^36^ Secondly, it can be maintained out of equilibrium in a closed reactor for days.^37^ Thirdly, working in water at pH 7, it is *a priori* compatible with other biochemical reactions.^38^ Lastly, it generates a variety of reaction-diffusion patterns such as traveling fronts, ^39^ waves^15^ and stationary patterns. ^33^ In a first series of experiments, the chemical subsystem encoded an autocatalytic loop that produces ssDNA species **A** — the node— in the presence of ssDNA **T**_AA_ —the template—, a polymerase and a nicking enzyme^14^ (Figure 1a).

In the mechanical subsystem, the bundles are constituted of stabilized microtubule filaments assembled together by attracting forces generated by the presence of a depletion agent^18^ (Figure 1b). The motors are clusters of kinesin-1 and thus can bind several microtubules at once.^*†*^ Such an active gel generates macroscopic flows that, depending on the concentration of motors and filaments, produce a diversity of microtubule morphologies:^6,7^ local contractions,^16^ corrugations, ^25^ chaotic flows,^18^ and global contractions.^22^ In the following, we demonstrate that, when mixed together, the two subsystems retain their ability to undergo, respectively, chemical and mechanical instabilities that generate spatiotemporal patterns. We further show that the strength of the active flow generated by the mechanical subsystem controls the mechanochemical coupling between the two subsystems (Figure 1d). Finally, we take advantage of this property to design materials that mimic the mechano-chemical transduction mechanism defining the polarity of the *C. elegans* embryo^5^ (Figure 1e).

To check whether the two subsystems remained functional when combined in an optimized buffer (Figure S1), we tested the propagation of a DNA front through and active gel undergoing local contractions. To do so, a solution containing all the components of the chemical and mechanical subsystems, except the strand **A**, were filled into a microchannel.^*\**^ An initial condition containing the same solution supplemented with **A** was injected on the left side of the channel (Figure 1c). We recorded the spatiotemporal dynamics of each subsystem by fluorescence microscopy thanks to the presence of a DNA intercalator that becomes fluorescent upon binding to double-stranded DNA and of fluorescently-labeled microtubules (SI Methods). In the chemical subsystem, we observed the propagation of a front of DNA fluorescence with constant velocity *v*_*c*_ = 20 *µ*m/min across the whole length of the active gel, *i*.*e. >* 1 cm (Figures 1d, 2a, S2 and Movie S1). Concomitantly, in the mechanical subsystem the microtubules contracted locally with a characteristic time *τ*_*m*_ = 50 min into aggregates with a typical size of 100 −500 *µ*m. The observation of a DNA front and microtubule aggregates is in agreement with previous reports for each subsystem taken independently. ^16,39^

**Figure 2.**
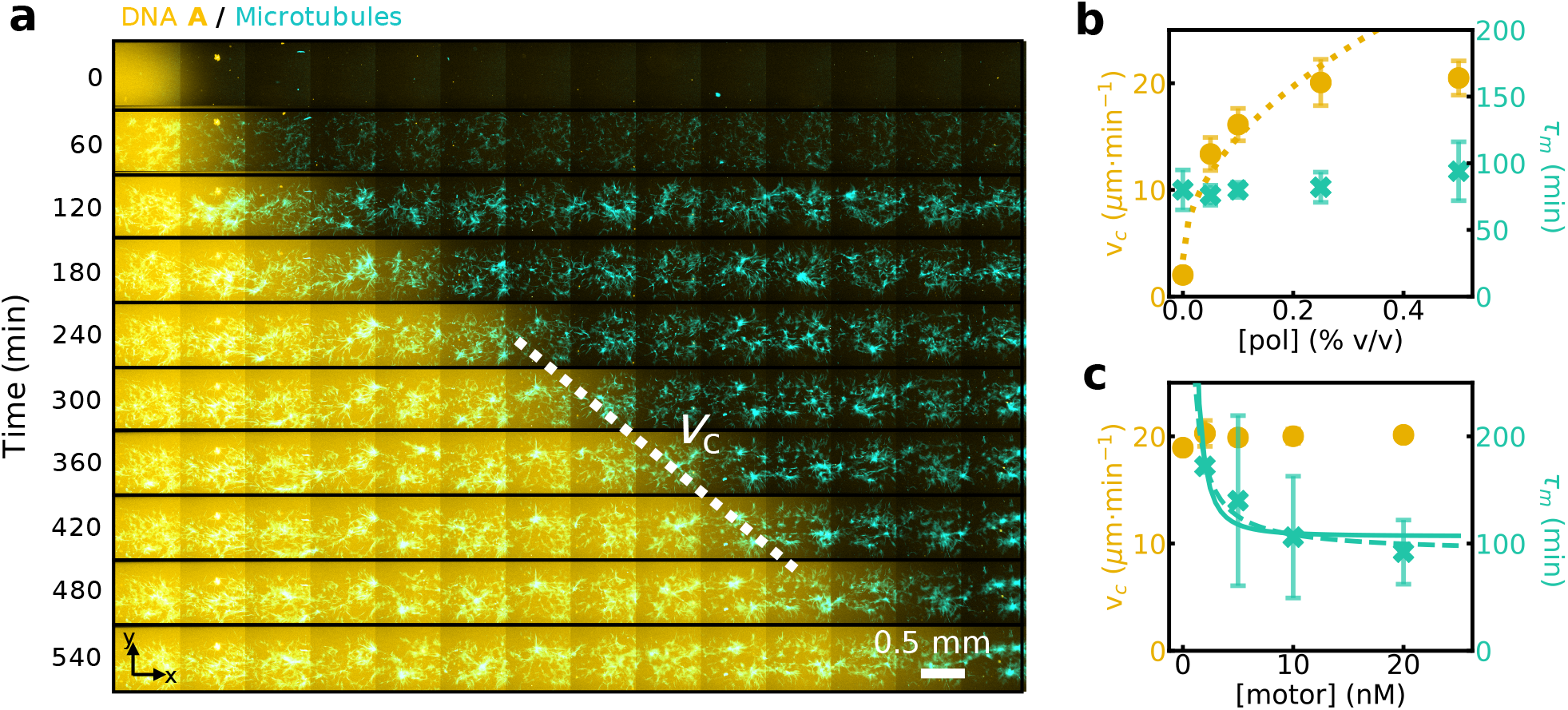
A DNA/enzyme reaction-diffusion front propagates normally inside a locally-contracting cytoskeletal active gel and the dynamics of each subsystem can be independently tuned. (a) Time-lapse 2-color image of the fluorescence intensities associated to species **A** (yellow) and to the microtubule network (light blue) (see also Movie S1). The dotted line indicates the velocity of the chemical front, *v*_*c*_. (b, c) Plots of *v*_*c*_ (yellow disks) and contraction time of the active gel, *τ*_*m*_, (blue crosses) for different concentrations of DNA polymerase (b) and motors (c). The lines are fits to the data with *v*_*c*_ ∼ [pol]^1*/*2^ (dotted line), *τ*_*m*_ ∼ [motor]^*−*1^ (dashed line) and *τ*_*m*_ ∼ [motor]^*−*2^ (plain line). Error bars correspond to one standard deviation from a triplicate experiment.

When the active gel produces local contractions, the dynamics of each subsystem can be independently tuned. Increasing the polymerase concentration, [pol], increases the front velocity, *v*_*c*_, until reaching a plateau at 22 *µ*m/min (Figure 2b). In these conditions, the characteristic contraction time *τ*_*m*_ remained constant. We find a scaling *v*_*c*_ ∼ [pol]^1*/*2^, in agreement with previous results^39^ and characteristic of Luther reaction-diffusion dynamics where 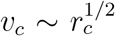, taking *r*_*c*_ ∼ [pol] for the rate of the autocatalytic reaction as observed in previous experiments.^39^ In turn, when the motor concentration, [motor], increases, *τ*_*m*_ decreases until reaching a plateau at 100 min (Figure 2c) and the size of microtubule aggregates increases (Figure S3), while *v*_*c*_ remains constant. A hydrodynamic model of a contracting active gel^7^ yields 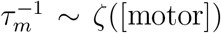, where *ζ*is the strength of the gel activity, which depends on the motor concentration, and two scalings are found in the literature ^19,40^ yielding *τ*_*m*_ ∼ [motor]^*−α*^, with *α* = 1 or 2 (Supplementary Text). Our data are compatible with both scalings (Figure 2c). In summary, Figure 2 shows that the two subsystems are both chemically and mechanically decoupled when the gel undergoes local contractions.

By varying the conditions, we can propagate the chemical front through active gels under-going other spatial instabilities associated with different active flow strengths, as sketched in Figure 1d. When the dGTP concentration was reduced (Figures S4-S5), chaotic flows were observed in the mechanical subsystem during several hours before local contractions occurred (Movie S2). Such flows did not modify the velocity of the chemical front because transport remained dominated by Brownian diffusion.^*\**^ When the length of microtubules was increased using taxol and the motor concentration reduced, the active gel formed corrugations reminiscent of those previously reported in the absence of the chemical subsystem,^25^ again without perturbing the chemical front (Movie S3).

In contrast, a dramatic perturbation of the front propagation was observed when the active gel underwent a global contraction, associated with large hydrodynamic flows (Figures 3 and S6-S11 and Movie S4). Global contractions were observed for long microtubules and relatively high motor concentrations (Figure S7). When the gel contracted more rapidly than the front propagated, the front moved faster and we distinguished 4 phases (Figure 3c). During phase I, the active gel contracted rapidly towards the center of the channel, accelerating until reaching a maximum velocity *v*_*m*_ = 400 *µ*m/min at the end of phase I and dragging DNA along, which formed a detached DNA islet ahead of the front. During phase II the gel decelerated and the DNA islet was diluted in the *xy* plane resulting in a front with a skewed profile 5-fold wider than the initial one (Figure S6). Throughout phases I and II the Pé clet number was greater than 1 (Figure S8), indicating that active convection predominated over diffusion. Finally, when the active gel stopped contracting, the DNA front slowly recovered a sigmoidal shape (phase III, Figure S6) and eventually reached a steady state with constant velocity and width (phase IV).

**Figure 3.**
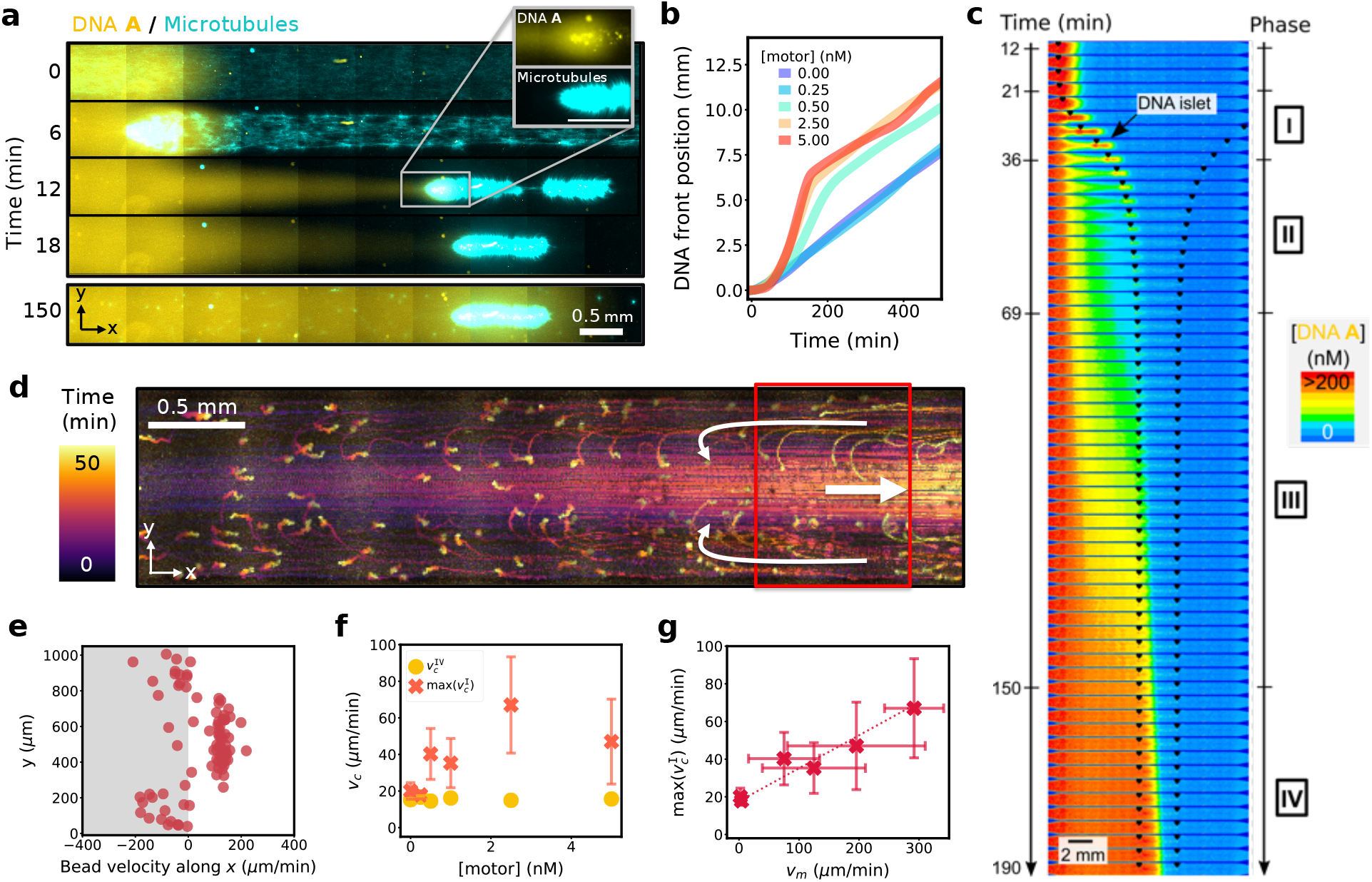
A globally-contracting active gel stretches and accelerates a reaction-diffusion front. (a) Time-lapse, 2-color fluorescence image with the DNA front in yellow and the microtubules in light blue. The inset shows the two fluorescence channels in separate images for the selected region. The white and yellow spots are dust particles concentrated by the contracting gel (see also Movie 4). (b) Position of the DNA front along *x* for different motor concentrations. (c) Time-lapse images of DNA fluorescence (color) at [motor] = 2.5 nM and 3 min per image. The extremities of the contracting gel are indicated with black markers. Roman numbers indicate the 4 phases described in the text. (d) Stroboscopic image averaged over 50 min showing the trajectories of fluorescent beads during gel contraction, the white arrows indicate the sense of the flow (see also Movie 5) and (e) plot of the bead velocity along *x* across the width of the channel for the beads in the red rectangle, 25 min after the beginning of the contraction. (f) Maximal front velocity during phase I (crosses) and steady-state velocity during phase IV (disks) for different motor concentrations. (g) Maximal front velocity during phase I *vs*. maximal gel contraction velocity. Error bars correspond to one standard deviation from a triplicate experiment.

In a control experiment with a passive dye, only phases I and II were observed, indicating that reaction was necessary for phases III and IV and that DNA islet formation was not related to the binding of DNA to the active gel (Figure S10). We confirmed the last interpretation by adding passive brownian beads to measure the hydrodynamic flow induced during contraction. We observed two counter-rotating fluid rolls, symmetric along the central axis of the channel, *x*, and producing water flows along *x* that reached +150 *µ*m/min in the center of the channel and −100 *µ*m/min at its borders (Figures 3d,e and S11 and Movie S5). We thus conclude that the stretching of the concentration profile of **A** leading to the formation of the DNA islet during phase I was a purely hydrodynamic process.

The active gel contraction velocity, *v*_*m*_, was a sigmoidal function of the motor concentration (Figure S7). We quantified the two main regimes of front propagation with max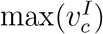 the maximal velocity during phase I and 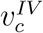 the velocity at steady state in phase IV. Figure 3f shows that the former strongly depended on the motor concentration while the latter was independent. Finally, the linear relationship between max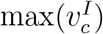 and *v*_*m*_ is consistent with the observation of a convection-dominated transport during phase I (Figure 3g). Taken together, these results show that when the flows generated by the active gel are sufficiently fast there is a mechano-chemical coupling between the gel and the reaction-diffusion front. This coupling happens through hydrodynamics and can be interpreted as a time-dependent Taylor dispersion. ^41^

We have just seen that active flows can significantly modify heterogeneous concentration profiles present in the chemical subsystem. In the following, we will show that homogeneous concentration profiles can also be mechanically modulated and we will engineer a mechano-chemical transduction pathway in this context. This is precisely what happens in the *C. elegans* embryo, where the active flow generated by the actomyosin cortex breaks the symmetry of an initially homogeneous distribution of PAR-proteins. Later, this asymmetry is amplified by a PAR-dependent bistable reaction network, leading to embryo polarization. ^5^ The reaction-diffusion active matter system developed here is a good candidate to mimic this process in a synthetic material. To do so, we first need to implement a mechanism that couples a variation in the microtubule concentration with a change in the concentration of a DNA species that is initially homogeneously distributed. Second, we need to engineer a chemical network that amplifies this concentration change.

The first requirement was fulfilled by attaching DNA strands to ∼ 30 *µ*m-diameter hydrogel beads, which were trapped by the microtubule mesh and concentrated during contraction. The second condition was satisfied by engineering a chemical subsystem whose kinetics depend on the concentration of DNA-bead conjugates, and thus on the contraction state of the gel. More precisely, the DNA autocatalytic loop was split into two nodes, **B** and **D**, that cross-activate each other thanks to the templates **T**_BD_ and **T**_DB_ (Figure 4a). By attaching each of these templates to a set of hydrogel beads and supplementing the medium with an exonuclease that degrades **B** and **D** (Figure 4a) the cross-catalysis kinetics become diffusion-controlled^42^ and thus should depend on bead density. As a result, the beads brought together in a contracted gel should activate faster, producing DNA that light them up in the presence of a DNA intercalator dye (Figure 4b). Indeed, when both types of beads where embedded in the active gel in the presence of a homogeneous, low concentration of **D**, they first reached a high density as the gel contracted and later they became fluorescent (Figures 4c,d, S13 and Movie S6). In the absence of contraction, the bead fluorescence amplification was delayed (Figure S12) and its final amplitude reduced, both by a factor 2 (Figure 4d). As expected for diffusion-controlled kinetics, the mechano-chemical DNA amplification dynamics slowed down with increasing exonuclease concentration (Figure 4d).

**Figure 4.**
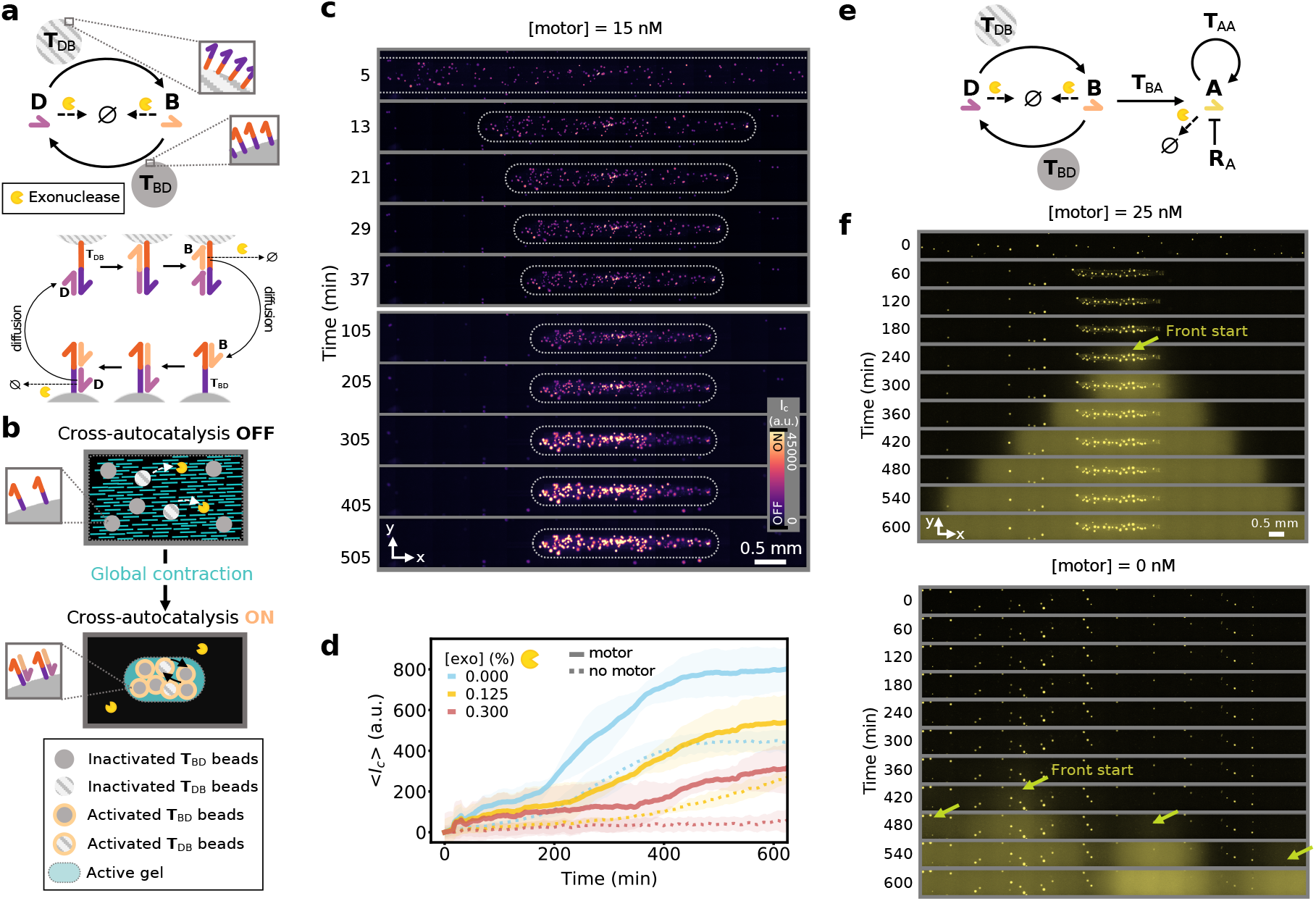
A globally-contracting active gel triggers the activation of downstream reaction networks with temporal and spatial control. (a) Scheme of the cross-autocatalytic DNA/enzyme network (top). Plain and dotted arrows indicate activation and degradation reactions, respectively. Harpoon-ended arrows correspond to ssDNA and disks indicate hydrogel beads carrying templates **T**_ij_. Detailed mechanism of bead activation where the diffusion of **B** and **D** between beads is indicated (bottom). (b) Sketch of the mechano-chemical activation of the reaction network in panel a through the contraction of the active gel (light blue) that brings the hydrogel beads (disks) close together, speeding up cross-catalysis. (c) Time-lapse images of DNA fluorescence from the template-bearing beads embedded in the active gel in the presence of motors. White dotted lines indicate the borders of the active gel and the channel walls are depicted in gray. (d) Average DNA fluorescence over the whole channel *vs*. time in the absence (dotted line) and in the presence (plain line) of motors for different exonuclease concentrations (colors). (e) Scheme of the bead-associated cross-autocatalytic network coupled to the autocatalysis of **A** in solution. Disks indicate templates linked to hydrogel beads. The blunt-ended arrow indicates repression. (f) Time-lapse images of DNA fluorescence in the channel for the network in panel e, in the presence (top) and in the absence (bottom) of motors. The bright spots are the beads, the arrows indicate the start of the fronts of **A**, in yellow.

To show that the activated beads may trigger downstream reactions in solution, the previous system was supplemented with freely-diffusing templates **T**_BA_ and **T**_AA_, that respectively convert **B** into **A** and sustain the autocatalytic reaction of **A** described earlier. In addition, to suppress the undesired self-activation of **T**_AA_, a repressor strand **R**_A_ was added^36^ (Figure S15). In the presence of motors, the beads were all activated within 1 h at the center of the channel, where the gel contracted, and they triggered a controlled front of **A** that propagated from the center of the channel to its extremities (Figures 4f and S14-S15 and Movie S7). In contrast, in the absence of gel contraction, the beads randomly activated over the course of 5h, which was followed by the uncontrolled amplification of **A**. Taken together, these results demonstrate that mechano-chemical transduction can be engineered to trigger either temporal or spatio-temporal chemical instabilities in a synthetic material.

The coupling of chemical and mechanical self-organization is a key ingredient of biological complexity, in particular during embryogenesis. We have demonstrated that it is possible to couple two archetypal examples of these mechanisms, reaction-diffusion and active matter, in a synthetic material. Our design is modular because it relies on two distinct subsystems with well-characterized and predictable spatiotemporal behaviors: DNA/enzyme reactions and kinesin/microtubule active gels. Considered independently, each subsystem reveals complex dynamics and macroscopic organizations which are subject to intense scrutiny.^15,16,18,20,22,25,33,38^ When mixed-together, the coupling strength between the two subsystems is set by the magnitude of the flow generated by the active gel. As a result, this system may be useful for investigating self-organization when chemical and mechanical out-of-equilibrium processes are intertwined. Finally, reaction-diffusion active matter provides a framework for the rational engineering of functional out-of-equilibrium materials with life-like properties. ^7,33,34^ On the one hand, it could be advantageously combined with the wide array of methods in DNA nanotechnology, such as nanostructure design,^43^ logic gates,^12^ analyte detection^44^ or hydrogel swelling. ^28^ On the other hand, by using DNA-motor conjugates^29–31^ or photosensitive motors^45^ the system is extendable to chemo-mechanical as well as photo-mechanical couplings.

## Acknowledgements

We thank H. Berthoumieux, V. Bormuth, M. Elez, A. Genot, G. Gines, N. Lobato-Dauzier, A. Maitra, L. Robert, Y. Rondelez and R. Voituriez for insightful discussions and K. Furuta and Z. Gueroui for their kind gift of kinesin plasmids. This work has been funded by the European Research Council (ERC) under the European’s Union Horizon 2020 programme (grant No 770940, A.E.-T.), by the Ville de Paris Emergences programme (Morphoart, A.E.-T.) and by MITI CNRS (J.-C. G.). The data that support the findings of this study are available from the corresponding authors upon reasonable request.

## Footnotes

† Two types of clusters were used: biotinylated kinesin-1 (from *D. melanogaster*) assembled together by streptavidin, ^16^ or clusters made from SNAPtag modified kinesin-1 (from *R. norvegicus*) that spontaneously multimerize ^25^ (SI Methods).

∗ The experimental conditions for each figure are provided in Tables S2-S5.

* The diffusivity of **A** due to the active flow, *D*_*f*_, was estimated to be 10-fold smaller than the Brownian diffusivity of **A**, *D*_*A*_, and thus 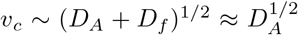 corresponds to a purely reaction-diffusion front (Supplementary Text).

