## Supplementary Information for "Programmed mechano-chemical coupling in reaction-diffusion active matter"

**5 Legends of Movies S1 to S7** **37**

**SI References** **39**

### 1 Materials and Methods

#### 1.1 Kinesin purification

Kinesins were purified as previously described (*1*). Two different kinesins were used: K430 and K401.

**K430.** From the plasmid coding for the K430 truncated kinesin-1 (amino acid residues 1-430) from Rattus norvegicus designed by Furuta et al. (*2*), we built a homodimer version containing a SNAP-tag on each arm. Furuta's plasmid, pET-32ark430(C7S)-FlagC-rk430(C7S)-SnapC-His, which expresses both Kinesin-Snap-His and Kinesin-Flag was digested at two EcoRI sites to split into two fragments of Kinesin-Flag sequence and Kinesin-Snap-His with backbone sequence coding ampicillin resistance gene. The digested linear sequences were purified by QIAquick PCR Purification Kit (Qiagen) and recircularized by T4 DNA ligase using Rapid DNA Ligation Kit (Thermo Scientific). The recircularized plasmids were transformed into DH5$\alpha$ competent cells and selected on an agar plate supplemented with 100 $\mu$g/mL ampicillin. The plasmids in colonies on agar plate were amplified in LB ampicillin medium and purified with Monarch Plasmid Miniprep Kit protocol (New England Biolabs). The sequence was verified by restriction enzyme digestion. Homodimer K430 was expressed in competent cells, Rosetta2 (DE3) (Novagen). The cells were cultured in LB supplemented with 100 $\mu$g/mL ampicilin and 34 $\mu$g/mL chloramphenicol at 37°C until OD660 = 0.6. The protein expression was induced by 0.1 mM IPTG for 5 hours at 22°C. The cells were collected and resuspended in the Lysis Buffer: Buffer A (20 mM Na-Pi buffer pH 7.5, 1 mM $MgSO_4$, 250 mM NaCl, 0.015 mM ATP, 10 mM $\beta$-mercaptoethanol, 0.1 % Tween-20) supplemented with 10 mM imidazole and 1X protease inhibitor cocktail (Sigma-Aldrich). The cell suspension was then sonicated (VCX-130, Sonics Materials) and centrifuged. The supernatant was collected and filtrated before mixing with Ni-IMAC resin (Biorad). The resin was washed first with the Buffer A supplemented with 10 mM imidazole, and then with the same buffer but with 50 mM of imidazole. The resin was eluted with the Buffer A supplemented with

250 mM imidazole. The eluate was filtrated and placed in a 14 kDa MWCO cellulose tube (Sigma-Aldrich) and then dialyzed with Buffer A three times (twice for 1 hour and then overnight) at 4°C. The dialyzed protein was further purified with a Superdex 200 Increase column (GE Healthcare). The kinesin corresponding peak fraction was collected, flash-frozen and kept at -80°C.

**K401.** For K401 purification, pT7-7_DmKinesin 1-401 BCCP-CHis6 (pWC2) plasmid designed by Gelles J. J (*3*) (Addgene plasmid # 15960 ; http://n2t.net/addgene:15960 ; RRID:Addgene_15960) was expressed in competent cells Rosetta2 (DE3) (Novagen). The cells were cultured in LB supplemented with 100 $\mu$M biotin, 100 $\mu$g/mL ampicillin and 34 $\mu$g/mL chloramphenicol at 37°C until OD660 = 0.7. The protein expression was induced by 1 mM IPTG for 2 hours at 22°C and then the biotinylation was induced by 0.2 mM rifampicin for 20 hours at 22 °C. The cells were collected and resuspended in the Buffer B (50 mM PIPES pH 7.2, 4 mM $MgCl_2$, 50 $\mu$M ATP, 10 mM $\beta$-mercaptoethanol) supplemented with 20 mM imidazole and 1X protease inhibitor cocktail (Sigma-Aldrich). The cell suspension was then sonicated (VCX-130, Sonics Materials) and centrifuged. The supernatant was collected and filtrated before mixing with Ni-IMAC resin (Biorad). The resin was washed with the Buffer B and 20 mM imidazole. The resin was eluted with the Buffer B and 500 mM imidazole. The eluate was placed in a Float-A-Lyzer G2 (5 mL; 50 kDa MWCO, Spectora/Por), dialyzed with Dialysis Buffer (250 mM PIPES pH 6.7, 20 mM $MgCl_2$, 0.25 mM ATP, 50 mM $\beta$-mercaptoethanol) three times (1 hour, 2.5 hours and overnight) at 4°C. The dialyzed protein supplemented with 36 % sucrose and 2 mM dithiothreitol (DTT) was flash-frozen and kept at -80°C.

#### 1.2 Microtubule polymerization

Tubulin and TRITC-labeled tubulin were purchased from Cytoskeleton, Inc., dissolved at 10 mg/mL in 1X PEM-KOH buffer (80 mM PIPES pH 6.8 (adjusted with KOH), 1 mM EGTA, 1 mM $MgCl_2$), flash-frozen and stored at -80°C.

Taxol-stabilized microtubules were prepared in a polymerization mix consisting in 1X PEM-KOH, 1 mM GTP, 10 %(w/v) glycerol and tubulin at 5 mg/mL (including 2.5 % of TRITC-labeled fluorescent tubulin). The mix was incubated at 37°C for 15 min. 20 $\mu$M of paclitaxel (referred in the MT and in the following as taxol) was added to the mix and let at 37°C for five more minutes. After polymerization, newly-formed microtubules were centrifuged at room temperature for 10 min at 12000 g to remove free tubulin monomers. The microtubules were redissolved into 1X PEM-KOH, 1 mM GTP, 10 % glycerol, 20 $\mu$M taxol and kept in the dark at room temperature for few days.

GMPCPP (purchased from Jena Bioscience) was also used to form microtubules. They were polymerized in the presence of 1X PEM-KOH, 0.6 mM GMPCPP, 0.2 mM DTT, 10 %(w/v) glycerol and tubulin at 5 mg/mL (including 2.5 % fluorescent tubulin), at 37°C for 30 min and left at room temperature for 5 hours. They were used within the same day or flash-frozen and stored at -80°C.

#### 1.3 Hydrogel beads functionalization

DNA-coated sepharose beads were prepared as previously described (*4*). Streptavidin sepharose beads (34 $\mu$m median diameter) were purchased from GE Healthcare and were supplied in a 20 % ethanol solution. The concentration of the stock solution was 16,000 beads/$\mu$L. We prepared 100 $\mu$L of solution containing 2500 beads/$\mu$L functionalized with 0.2 nmol of biotin-DNA for these $10^5$ beads, at room temperature for 20 min in the Buffer B with a gentle agitation. Buffer B is composed of 0.6X PEM-KOH, 0.2X PEM-NaOH (for 1X: 80 mM PIPES, 1 mM EGTA, 1 mM $MgCl_2$ adjusted with 135 mM NaOH to pH 7.1), 8.0 mM $MgSO_4$ and 1.5 %(w/v) Pluronic F127. The beads were then subjected to three cycles of centrifugation (10,000 g, 30 s), removal of supernatant and wash with Buffer B. DNA-coated sepharose beads were finally conserved for few weeks at 4°C.

#### 1.4 Preparation of solutions

Solutions were prepared from the Buffer AM and the components listed in Table 2-Table 5. Buffer AM is composed of 0.6X PEM-KOH (for 1X: 80 mM PIPES, 1 mM EGTA, 1 mM $MgCl_2$ adjusted with 130 mM KOH to pH 6.9), 0.2X PEM-NaOH (for 1X: 80 mM PIPES, 1 mM EGTA, 1 mM $MgCl_2$ adjusted with 135 mM NaOH to pH 7.1), 8.0 mM $MgSO_4$, 1.5%(w/v) Pluronic F127, 1.0 mM ATP, 10 mM creatine phosphate, 5.0 $\mu$g/mL creatine kinase, 3.0 mM DTT, 20 mM D-glucose, 1.0 mM Trolox, 0.5 mg/mL BSA, 0.5X SYBR Green I. Glucose oxidase and catalase were not included in experiments involving DNA attached on hydrogel beads because we found that it caused variation in their fluorescence intensity.

The three enzymes needed for the chemical subsystem were: the Bst DNA polymerase Large Fragment (New England BioLabs) – abbreviated *Bst LF* or *pol* –, the nicking enzyme Nt.BstNBI (New England BioLabs) – abbreviated *NBI*. The exonuclease ttRecJ was produced in-house as previously described (*5*). The DNA sequences used in this work are listed in Table 1 and were purchased from Biomers.

Experiments were performed within 18 mm×1 mm×0.2 mm PMMA channels (Fluidic 152, ChipShop). For front propagation experiments, they were filled with 4.5 $\mu$L of the solution containing all the elements except the DNA **A**. The initial condition – containing the same solution with 1 $\mu$M of the DNA **A** – was injected on one side of the channel (0.25 $\mu$L). The surplus of liquid was carefully absorbed using a clean tissue. Finally, both ends of the channel were closed using vacuum grease, starting from the side without the DNA **A** to avoid contamination and the appearance of a second front that would propagate in opposite direction to the main front.

#### 1.5 Imaging

Epifluorescence images were obtained with a Zeiss Observer 7 automated microscope equipped with a Hamamatsu C9100-02 camera, a 10X or a 4X objective, a motorized stage and controlled with MicroManager 1.4. Images were recorded automatically every 3 s to 8 min (depending on the experiment) using an excitation at 470 nm (observation of the DNA intercalator) and/or 550 nm (observation of the microtubules) with a CoolLED pE2 or a CoolLED pE-4000. The temperature was controlled thanks to a transparent TokaiHit ThermoPlate. For optimal thermal conduction, mineral oil was added between the PMMA channels and the thermoplate on the microscope. The set-up was put in place at least 30 min before the filling of the channels in order to be sure that the temperature equilibrium (25°C or 28°C) was reached.

#### 1.6 Autocatalytic network $\mathbf{T_{AA}}$ dynamics in well-mixed tubes

Optimization experiments presented in Figure S1 were performed in well-mixed 200 $\mu$L tubes in a BioRad CFX qPCR machine using 10 $\mu$L of solution at the temperature of 25°C. Total DNA concentration was measured thanks to the intercalating molecule SYBR Green I (ThermoFisher Scientific) or Evagreen (Biotium).

#### 1.7 Data processing

Time-lapse images from microscopy experiments were processed by ImageJ/Fiji (NIH) and Python. Images of the channels were first cropped and rotated manually in order to compute the velocity of the chemical front $v_c$ and the time related to the dynamics of the active gel $\tau_m$.

##### 1.7.1 Determination of $v_c$

The DNA front images were recorded using a 470 nm excitation. Using a Python routine, images were filtered using a gaussian filter. The intensity was normalized between 0 and 1. Finally, the contrast was adjusted by putting 10% of the highest (respectively lowest) intensity values to 1 (respectively 0). At each time the intensity was averaged along the width of the channel ($y$ axis). We fitted this 1D array of averaged intensity by the expression $I_c = \frac{e^{(x_c - x)/l_0}}{1+e^{(x_c-x)/l_0}}$ where $x_c$ is the position of the front (the intensity is equal to 0.5) and $l_0$ its width. Thus we obtained the position of the front as a function of time. We computed the velocity of the front by fitting this function by an affine fit. For the experiments presented in Figure 3, the velocity $\max(v_c^{\mathrm{I}})$ was obtained by first deriving the position and then by taking the first 5 maximum velocities. $v_c^{IV}$ was computed using a linear fit after the transient regime.

##### 1.7.2 Determination of $\tau_m$

Fluorescent microtubules were recorded using a 550 nm excitation. In order to quantify the dynamics of active structures we first applied a gaussian filter, normalized the intensity and then computed the standard deviation of the normalized intensity along the $y$ axis (width) for each position $x$ of the channel length and for each time point. The standard deviation was then normalized and we computed $\tau_m$ as the time needed to reach 0.5 of this normalized standard deviation.

##### 1.7.3 Determination of $v_m$

Active gel dynamics was obtained by manually tracking small aggregates and/or dust embedded in the active gel. We chose the aggregates to be close to the border on the left (where the DNA front was initiated). The velocities were computed as the time derivative of the position. $\max(v_m)$ was obtained by taking the average of the 5 maximal velocities.

##### 1.7.4 Tracking of beads

In the experiments of Figure 3d-e, 1 $\mu$m-diameter fluorescent beads (Estapor Fluorescent Functionalized Microspheres F1-XC 100, Merck) were added to the solution to study the velocity of the active gel and the surrounding fluid. The beads were used at the concentration of 0.0015 %(v/v). The tracking of the beads was performed using the ImageJ plugin *TrackMate* (*6*) that calculated trajectories which were then analyzed by a Python routine to compute the velocities (derivative of the positions).

### 2 Supplementary Text

#### 2.1 Finding experimental conditions compatible with both subsystems

##### 2.1.1 Temperature

The working temperature of the DNA/enzyme chemical subsystem is usually set according to the melting temperature of the DNA strands as well as to the optimum temperature of the polymerase and the nicking enzyme, and possibly the exonuclease. This gives a range of temperatures between 40°C and 50°C (*7*). More recently 37°C was achieved in order to reach physiological conditions (*8*). However, the activity and the lifetime of the active gel is greatly reduced as the temperature increases, hypothetically due to proteins denaturation and the formation of pluronic micelles that may lead to a decrease in activity (*9*). We thus built an autocatalytic system that was able to work at 25°C. This system is based on a previous design that used the nicking enzyme Nt.BstNBI (New England BioLabs) (*10*). We shortened the node strand (**A**) from 11 to 10 nucleotides and found that this was sufficient to obtain an autocatalyst with a typical amplification time $\tau_{c,1/2} = 100$ min at 25°C (Figure S1 and (*11*)).

##### 2.1.2 Buffer

Besides the temperature, the buffer needed to be adjusted. Starting from the canonical buffer for experiments involving microtubules and kinesin motors (the PEM-KOH buffer also known as the BRB80 buffer (*12*)), we created a modified version by incorporating sodium cations (via the buffer PEM-NaOH, a PEM buffer adjusted in pH with NaOH) and magnesium cations (see section 1.4 for the composition). To assess conditions that would not be too deleterious for the active gel we tested the effect of the PEM-NaOH buffer and the magnesium ions.

First, we set the maximum concentration of the PEM buffers to 0.75X and prepared a mix between PEM-NaOH and PEM-KOH buffers. Thus, we show in Figure S1 (a) and (b) that sodium ions accelerated the autocatalytic amplification compared to potassium ions. We finally choose a mixture of PEM-NaOH at 0.2X and PEM-KOH at 0.6X. Such a change represented a gain in time of about 30% compared to a mix that only contained the PEM-KOH buffer.

Secondly, we added magnesium cations to the buffer (Figure S1 (c) and (d)). The presence of at least 10 mM $Mg^{2+}$ is needed for the DNA/enzyme chemical subsystem activity — in the form of $MgSO_4$ or $MgCl_2$ — compared to previous experiments involving the same system (*13*, *14*). It is hypothetically due to the presence of creatine phosphate which might be able to capture magnesium ions. We chose to add 8 mM of $MgSO_4$ (in addition to the 1.6 mM $MgCl_2$ already existing in the PEM buffers.

We did not notice any changes on the active gel activity when using this new buffer.

#### 2.2 Details on the model of a contractile gel

To estimate the characteristic time of local contractions, $\tau_m$, we model the active gel using a hydrodynamic equation supplemented with an active stress term (*15*). We consider a 1D active gel of length $L$ undergoing a constant active tension $\zeta\Delta_r G$ generated by the motors along the $x$ coordinate, with $\zeta$ the strength of the gel activity and $\Delta_r G$ the free energy associated to ATP hydrolysis by the motors. The force balance equation neglecting inertia is

$$0 = \eta \frac{d^2 v_m}{dx^2} + \frac{d\zeta\Delta_r G}{dx}, \tag{1}$$

where $\eta$ is the viscosity of the active gel and $v_m$ its velocity. The last term cancels out because the active tension is uniform. We thus have $0 = \eta \frac{d^2 v_m}{dx^2}$. Integrating this last equation, we find that the initial velocity of contraction is $v_m = -\frac{\zeta\Delta_r G}{\eta} x$.

We compute $\tau_m^{-1} = v_m/x = \zeta\Delta_r G/\eta$ and thus $\tau_m^{-1} \sim \zeta$ as indicated in the Main Text. $\zeta$ depends on the motor concentration and two scalings are found in the literature: $\zeta \sim [\text{motor}]^2$ in a 2-dimensional active gel similar in composition to ours but locally-extensile (*16*) and $\zeta \sim [\text{motor}]$ in a 3-dimensional globally-contracting gel in *Xenopus* oocyte extracts driven by dyneins (*17*).

#### 2.3 Hypothesis on the origin of the contractile behavior in presence of dGTP

dGTP dramatically changed the behavior of the active gel, which did not happen with the three other dNTPs (Figures S4 and S5). To the best of our knowledge this behavior has not been described and here we only formulate hypotheses that could explain it. Firstly, a nucleotide exchange may occur between GMPCPP and dGTP, as it happens with GTP (*18*), which would promote microtubule polymerization (*19*), and thus contraction. Moreover, the presence of dGTP leads to a microtubule growth-rate similar to the one in the presence of GTP (*18*, *19*). Secondly, the dynamic instability of microtubules — i.e. the stochastic switching between growth and shrinkage — may also play a role. Indeed, GMPCPP is known to suppress dynamic instability (*20*), contrary to dGTP (*18*). Lastly, it has been reported that microtubule binding proteins, such as kinesins, can discriminate between the nucleotide states of microtubules by having different affinities (*21*).

#### 2.4 Estimation of $D_f$

In the conditions of Movie S2 the typical velocity of the microtubule flow was estimated to be $v_m \approx 8$ $\mu$m/min by tracking bright spots transiently appearing on the bundles. One vortex of bundles has a typical diameter $l_m \approx 125$ $\mu$m. We estimate the dispersion coefficient of a passive tracer in the active flow as $D_f = l_m v_m = 1000$ $\mu\text{m}^2$/min. Zadorin et al.(*13*) measured the Brownian diffusivity of a single stranded DNA similar to **A** (11 nucleotides long instead of 10 as used here) and found $D_A = 12000$ $\mu\text{m}^2$/min at 25 °C. Thus $D_A/D_f \approx 12$ and $D_A \gg D_f$ indicating that the Brownian diffusion of **A** dominates over the dispersion induced by the chaotic flow.

### 3 Supplementary Figures

The experimental conditions for each figure are detailed in Tables S3-S4.

#### 3.1 Figure S1: Influence of the buffer composition on the autocatalytic network $T_{AA}$ dynamics in a well-mixed system

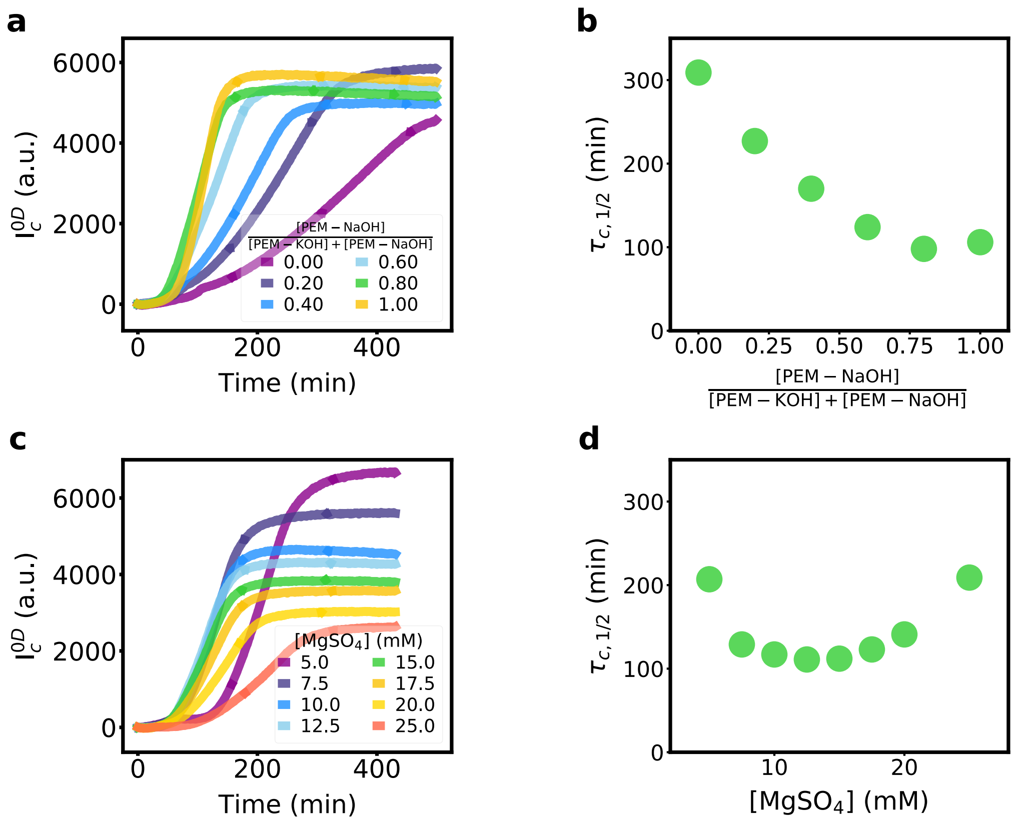

Figure S1: Influence of the buffer composition on the autocatalytic network dynamics in a well-mixed system. (a) Dynamics of the $\mathbf{T}_{AA}$ autocatalyst and (b) evolution of the time $\tau_{c,1/2}$ (time needed to reach half of the maximum intensity value) as a function of the ratio $\frac{[\text{PEM-NaOH}]}{[\text{PEM-KOH}]+[\text{PEM-NaOH}]}$. The total concentration of PEM buffers was set to 0.75X. This experiment was performed in the indicated PEM buffer supplemented with 6 mM $MgSO_4$ and supplemented with 1X Evagreen, 0.5 mM of each dNTP, 0.2 %(v/v) BstLF, 2.5 %(v/v) NBI, 200 nM $\mathbf{T}_{AA}$ and 10 nM **A**. (c) Dynamics of the $\mathbf{T}_{AA}$ autocatalyst and (d) evolution of the time $\tau_{c,1/2}$ (time needed to reach half of the maximum intensity value) as a function of $MgSO_4$ concentration. This experiment was performed in 0.15X PEM-KOH, 0.60X PEM-NaOH, 1X Evagreen, 0.5 mM of each dNTP, 0.2 %(v/v) BstLF, 2.5 %(v/v) NBI, 200 nM $\mathbf{T}_{AA}$ and 10 nM **A**. Both experiments were performed at 25°C.

##### 3.2 Figure S2: Fluorescence intensity related to the concentration of A over time at different positions along the channel during the propagation of the DNA/enzyme reaction-diffusion front (SI for Figure 2)

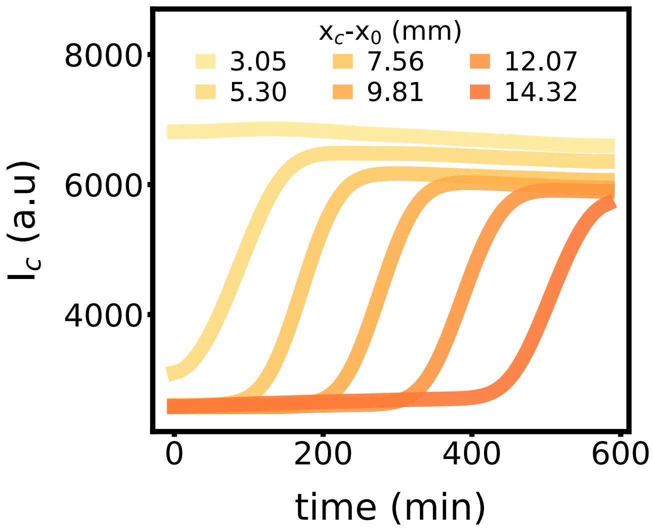

Figure S2: Fluorescence intensity related to the concentration of **A** over time at different positions along the channel during the propagation of the DNA/enzyme reaction-diffusion front in a locally-contracting active gel. Figure associated to Figure 2a in the Main Text.

##### 3.3 Figure S3: Morphology of the active gel after 300 min as a function of motor concentration (SI for Figure 2)

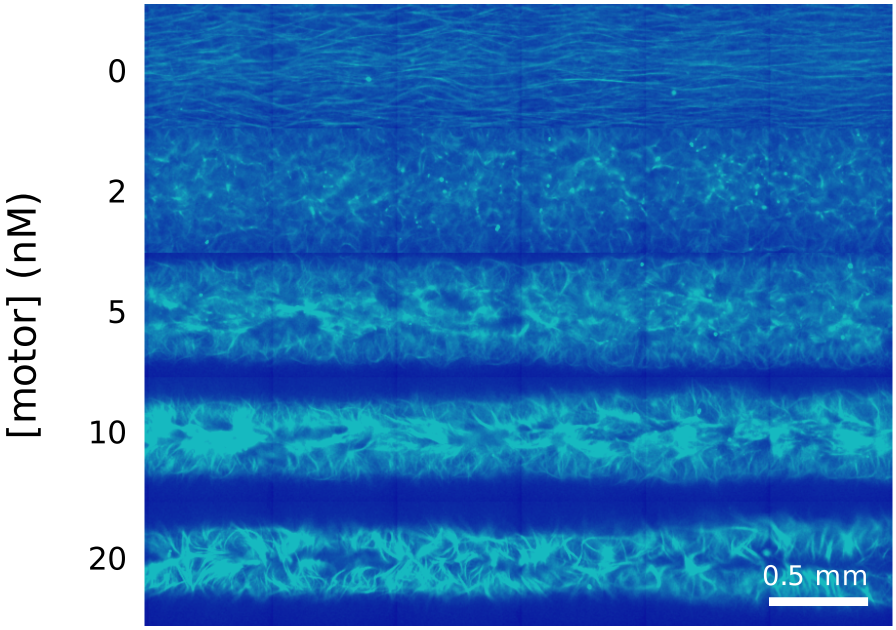

Figure S3: Morphology of the active gel after 300 min as a function of motor concentration after a DNA front of **A** has propagated across the gel (not shown). Microtubules were polymerized with GMPCPP. Data associated to Figure 2c in the Main Text.

#### 3.4 Figure S4: dGTP is responsible of the contractile behavior of an active fluid composed of GMPCPP-microtubules

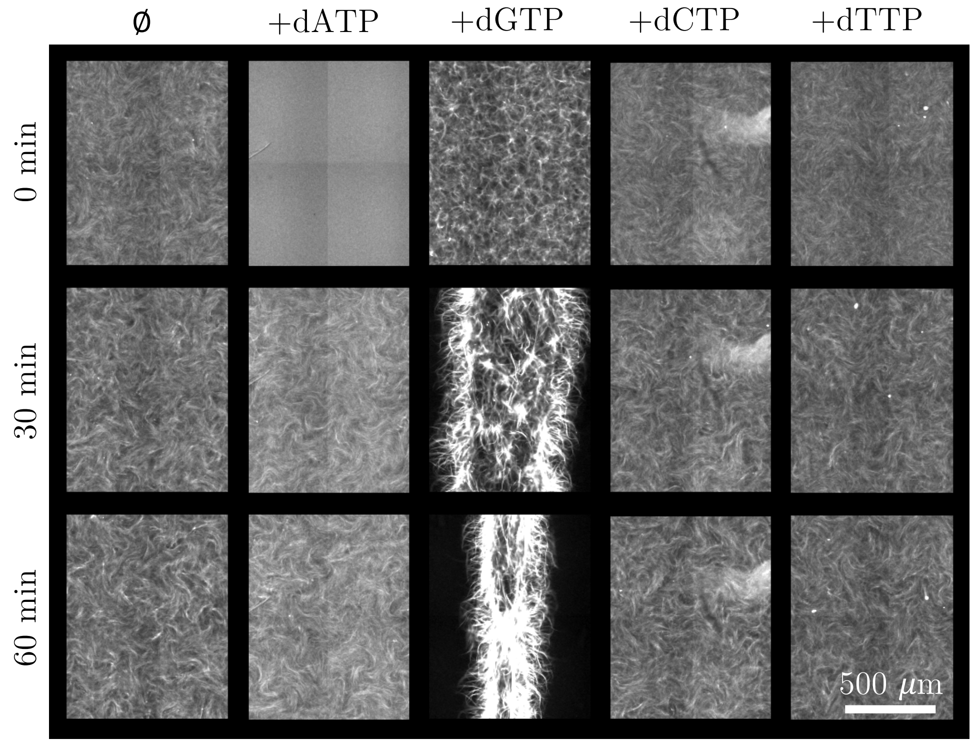

Figure S 4: dGTP is responsible for the contractile behavior of an active fluid composed of GMPCPP-microtubules. In the absence of the chemical subsystem, a GMPCPP-microtubules based active fluid is known to display chaotic flows (*22*). The presence of 0.5 mM of dGTP led to the contraction of the active network contrary to what was observed for the other dNTP.

##### 3.5 Figure S5: dGTP changes both the morphology of the active gel and tunes the velocity of the DNA front

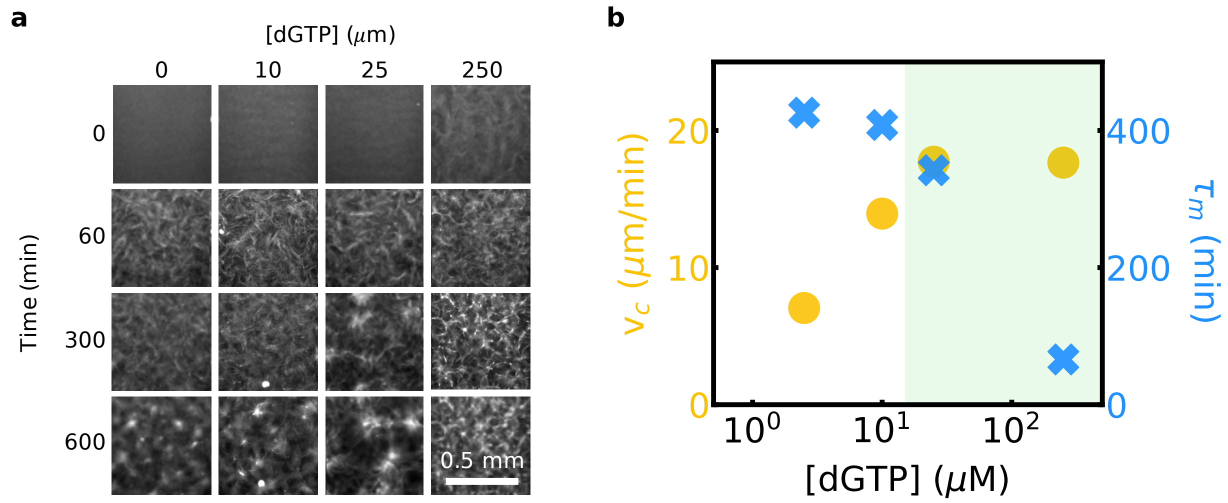

Figure S5: dGTP changes both the morphology of the active fluid and tunes the velocity of the front. (a) Images of the structures formed by the fluorescent microtubules at different times for different dGTP concentrations in the presence of the propagating DNA front. For [dGTP] <10 $\mu$M the active fluid displayed chaotic flows similar to those observed in the absence of the chemical subsystem (*22*). At higher dGTP concentrations, local contractions were observed after an initial chaotic flow regime whose lifetime was set by the dGTP concentration. (b) Velocity of the front (yellow disks) and contraction time of the active fluid, $\tau_m$ (blue crosses) as a function of dGTP concentration. The background color green indicates that the active fluid is locally contracting at least 60 min before the negative control ([dGTP] = 0 $\mu$M).

#### 3.6 Figure S6: Position of the DNA front over time for different motor concentrations (SI for Figure 3)

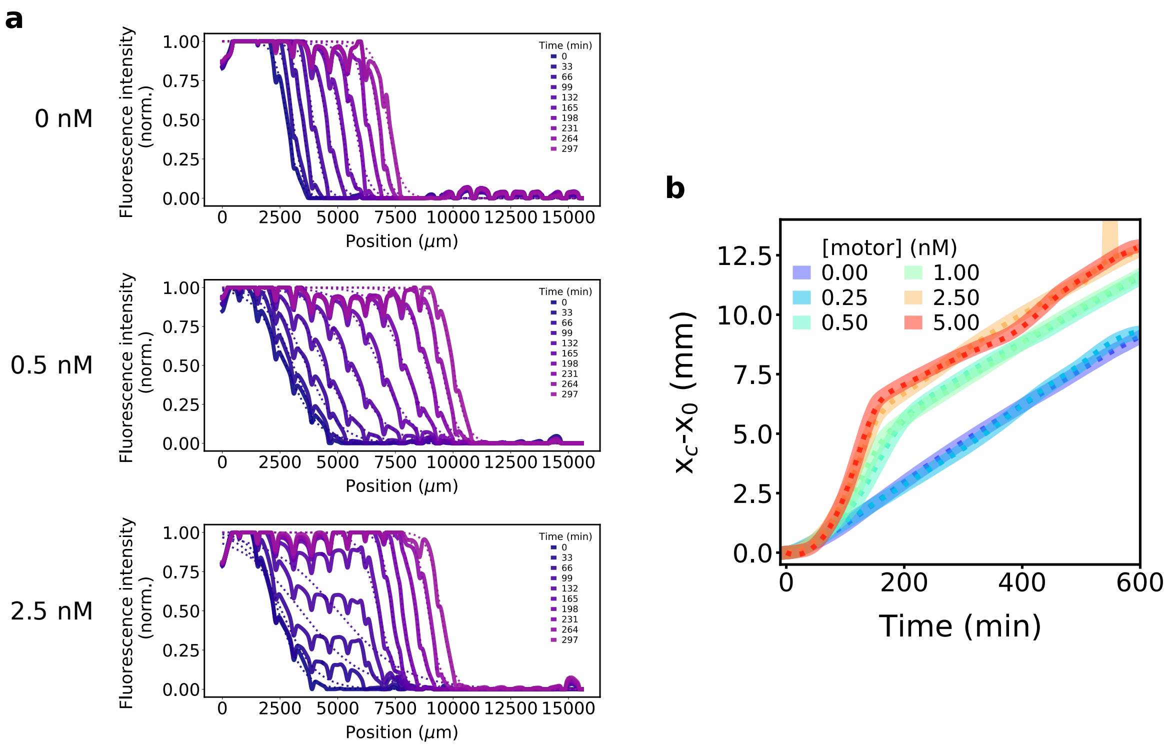

Figure S6: Position of the DNA front over time for different motor concentrations in a front propagating within a globally-contracting active gel. (a) Fluorescence intensity profiles of the DNA front for different motor concentrations and different times and their respective sigmoidal fit (dotted lines). (b) Evolution of the DNA front position $x_c(t)$-$x_0$ calculated from the sigmoidal fits of the fluorescence profiles.

##### 3.7 Figure S7: Active gel contraction dynamics (SI for Figure 3)

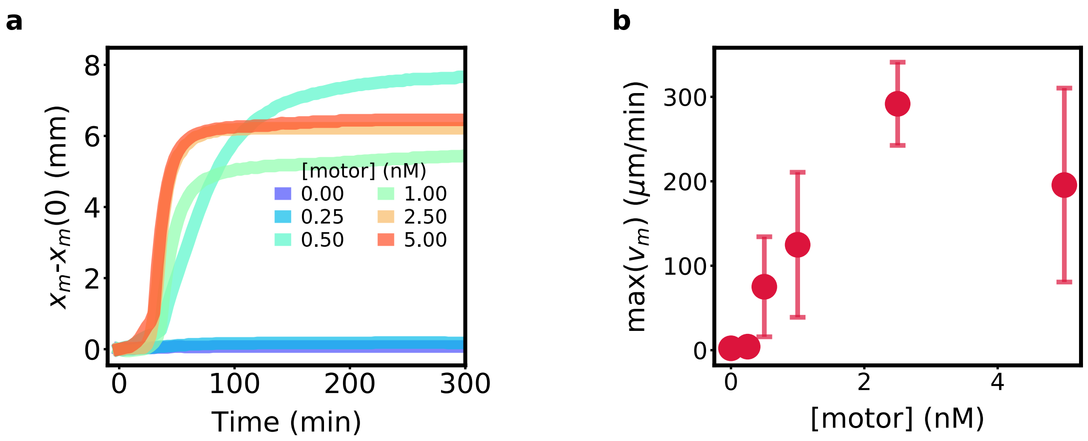

Figure S7: Dynamics of global contraction in the active gel. (a) Evolution of the gel position $x_m$-$x_m(0)$ (extremity on the left of the active gel) over time for different motor concentrations. (b) Maximal velocity of one border of the active gel max($v_m$) along the main axis of the channel (x-axis) as the function of kinesin motor concentration. Error bars correspond to one standard deviation from a triplicate experiment.

##### 3.8 Figure S8: Péclet number over time for different motor concentrations (SI for Figure 3)

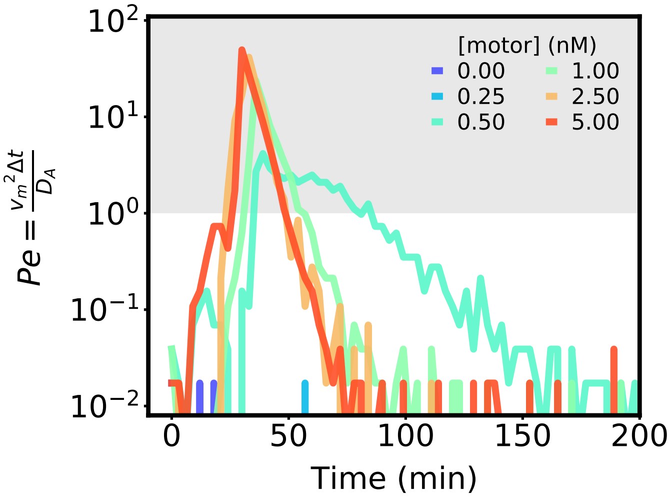

Figure S8: Péclet number over time for different motor concentrations. The gray region indicates a Péclet number superior to 1. We used $\Delta t = 3$ min and $D_A = 12000$ $\mu$m$^2$/min at 25 °C, a value found by Zadorin et al.(*13*) for a single stranded DNA similar to **A** (11 nucleotides long instead of 10 as used here).

##### 3.9 Figure S9: Influence of the microtubule concentration on $v_c$ and the contractile behavior (SI for Figure 3)

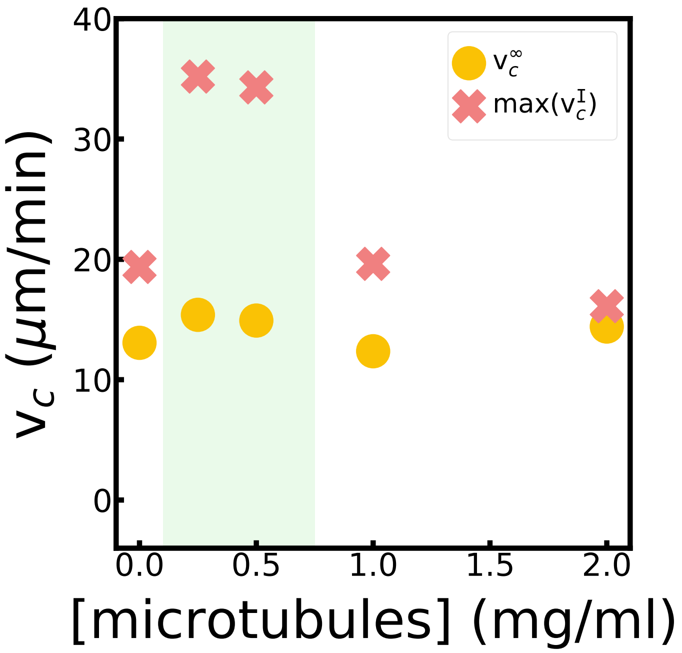

Figure S9: Influence of the microtubule concentration on $v_c$ and the contractile behavior. The green region indicates that the active gel was globally contracting. At high microtubule concentration, the active network was not anymore able to contract.

#### 3.10 Figure S10: Dilution and diffusion of fluorescein by the active gel (SI for Figure 3)

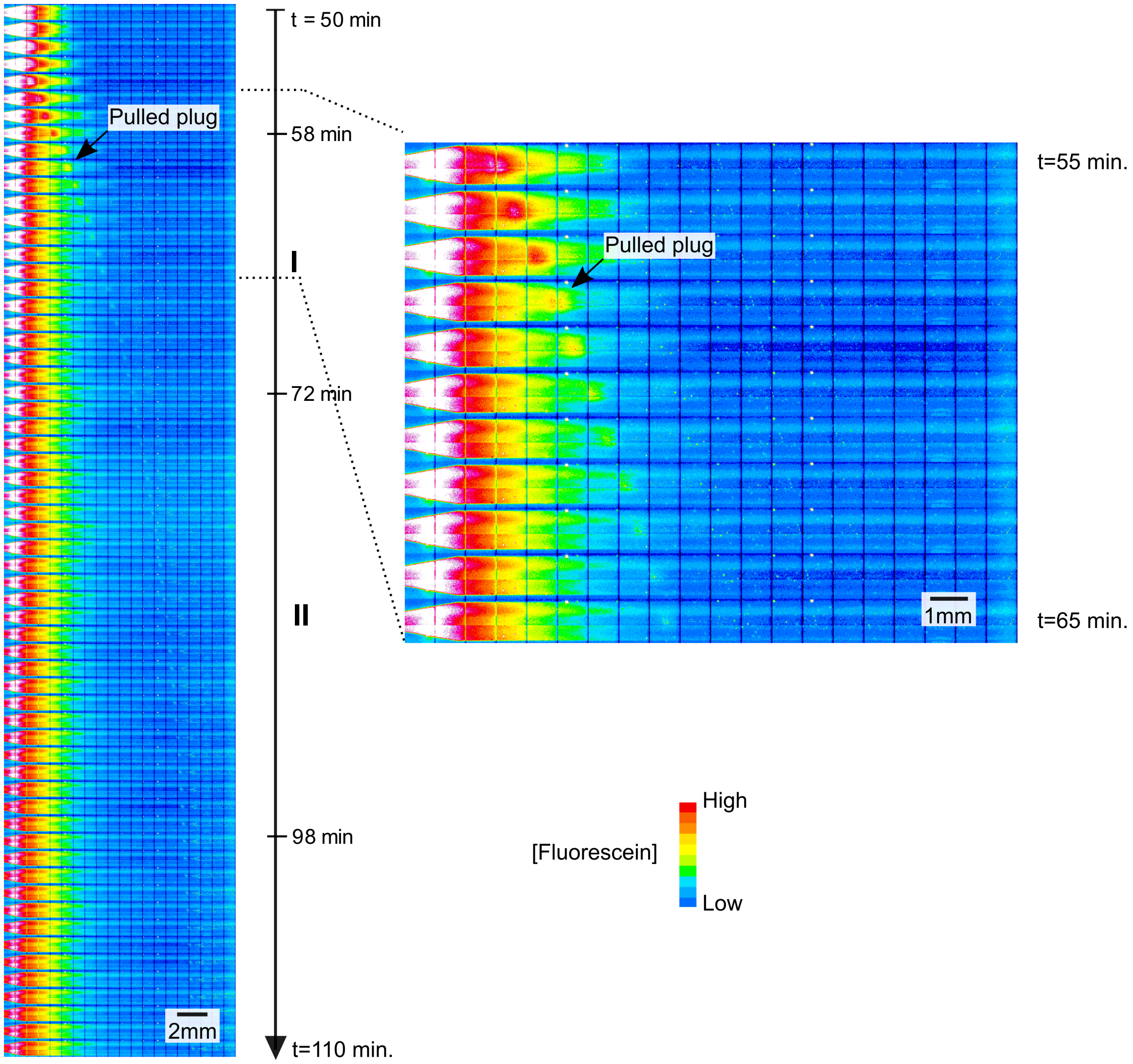

Figure S10: Dilution and diffusion of fluorescein by a globally-contracting active gel. Time-lapse image showing the position and shape of a 2 $\mu$M fluorescein plug introduced on the left side of the channel and perturbed by a globally contractile active gel. A 10-color lookup table was used to visualize the fluorescent islet pulled by the active gel. This mechanical phase (phase I) – similar to the one observed with DNA (see Figure 3 (c)) – indicated that the stretching step did not involve the binding between DNA and the active gel. During phase II the gel decelerated and the fluorescein islet (noted pulled plug here) was diluted by diffusion. The active gel was visible thanks to a small dust embedded in its left extremity (visible as a small light blue region from phase II). Note that in this experiment the active gel detached from the right border of the channel and then continued to contract for a longer time than in Figure 3c.

#### 3.11 Figure S11: Tracking of 1 $\mu$m-fluorescent beads at different positions and times during the global contraction of the active gel (SI for Figure 3)

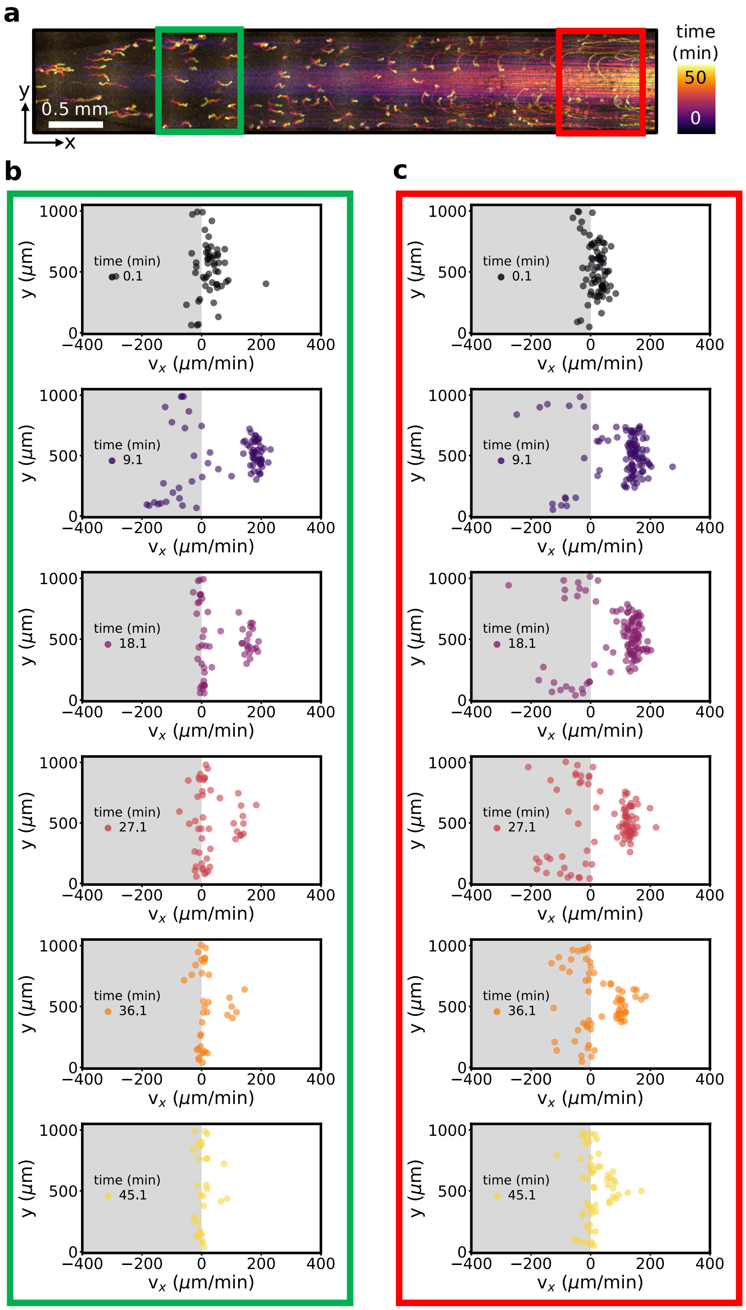

Figure S11: Tracking of 1 $\mu$m-fluorescent beads at different positions and times during the global contraction of the active gel. (a) Stroboscopic image of fluorescent beads during fluid contraction at [motor] = 20 nM. (b) and (c) Velocity along $x$ at different times for the fluorescent beads in the green and the red regions in panel a.

#### 3.12 Figure S12: In the absence of contraction the cross-autocatalysis was delayed and had a reduced amplitude (SI for Figure 4)

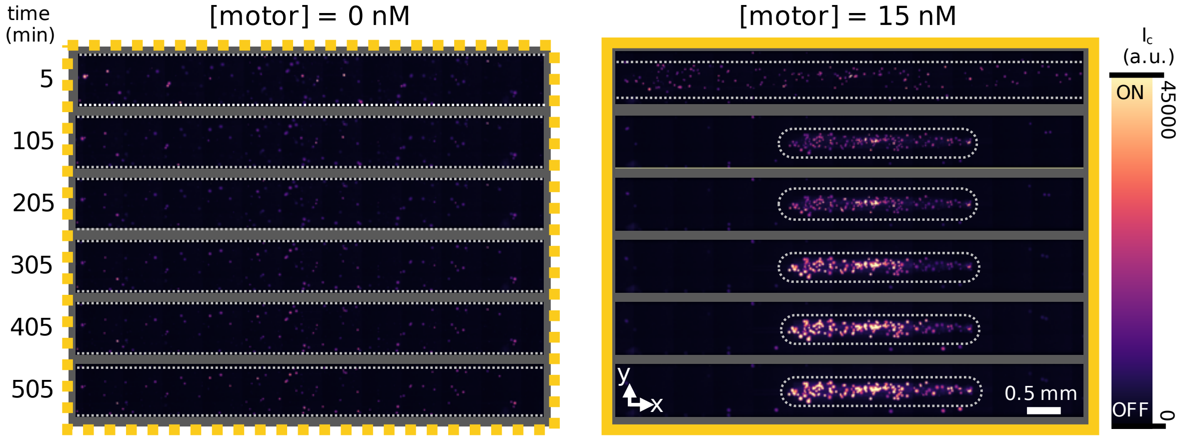

Figure S12: In the absence of contraction the cross-autocatalysis was delayed and had a reduced amplitude. Negative control (left) for Figure 4 (c) (data reproduced on the right panel).

#### 3.13 Figure S13: Local activation of the cross-autocatalytic network on beads (SI for Figure 4)

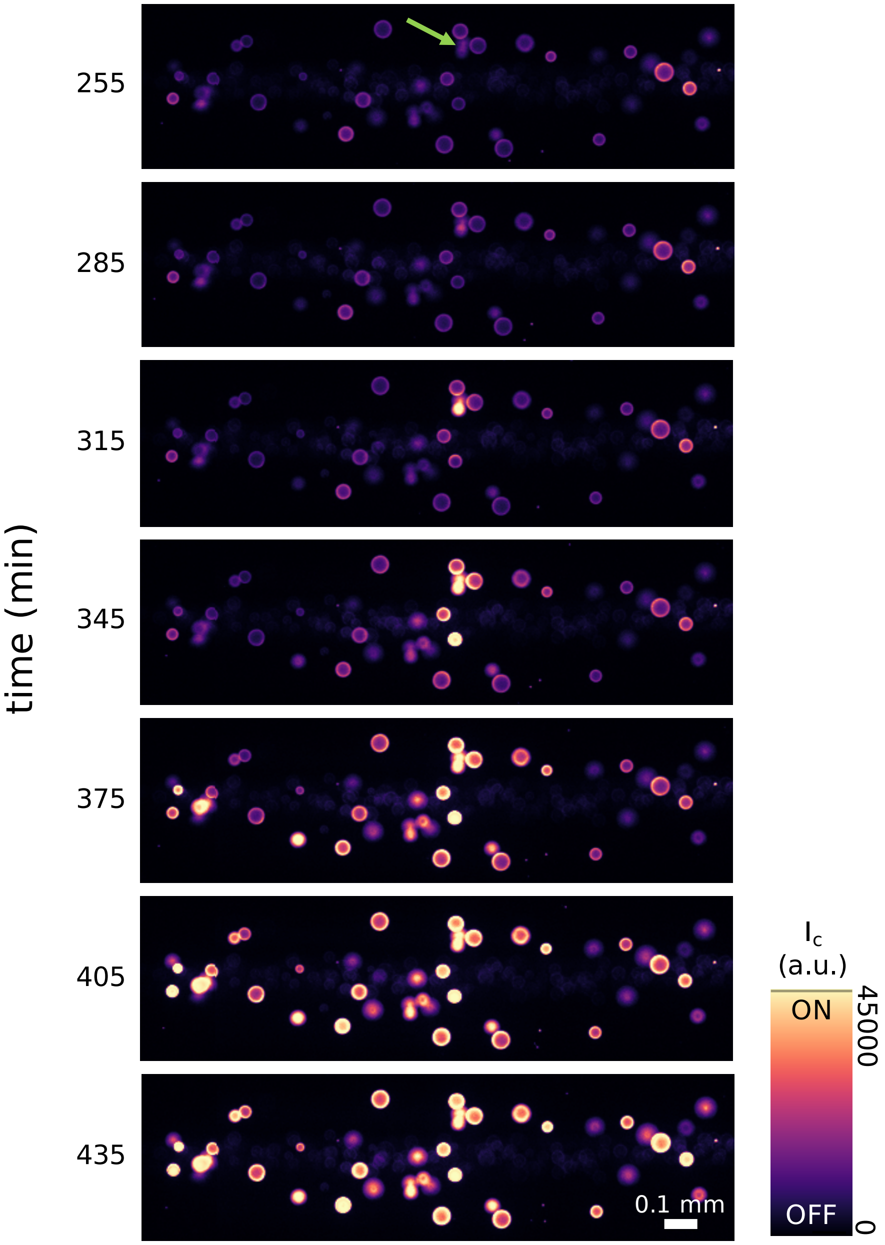

Figure S13: Time-lapse fluorescence images showing the activation of the cross-autocatalytic network when the beads are close to each other (after the contraction of the active gel). Because the cross-autocatalytic network was triggered by the DNA strand **D**, the $\mathbf{T}_{\mathrm{DB}}$ beads (green arrow) are the first to be active and then interact anisotropically (*4*) with the neighboring $\mathbf{T}_{\mathrm{BD}}$ beads.

#### 3.14 Figure S14: Influence of beads-$T_{DB}$ concentration on the triggering of a front of A (SI for Figure 4)

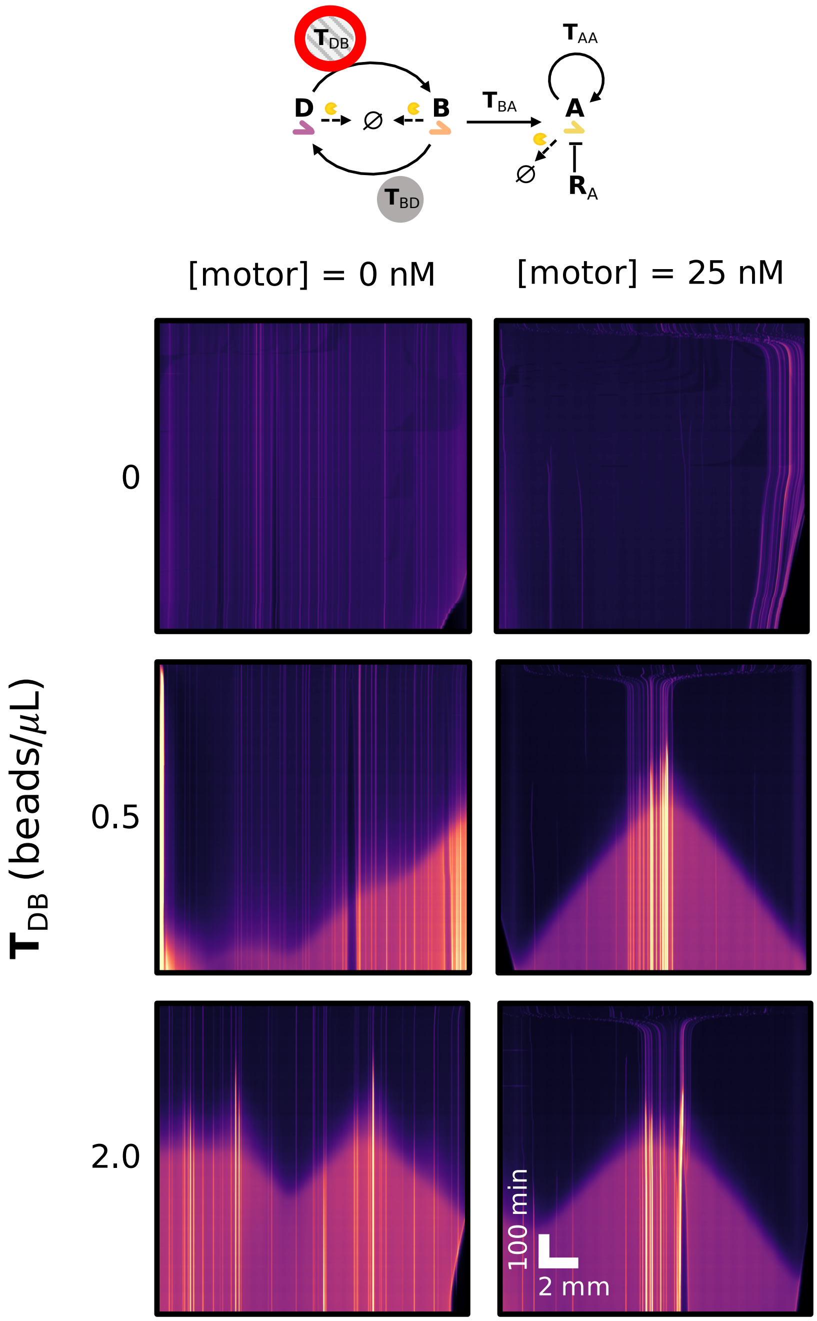

Figure S14: Influence of beads-$\mathbf{T}_{DB}$ concentration on the triggering of a front of **A**. Kymographs showing the activation of the beads and the propagation of a front of **A**. In the absence of the beads-$\mathbf{T}_{DB}$ we observed neither cross-autocatalytic amplification nor front. Increasing beads-$\mathbf{T}_{DB}$ concentration led to more activation and then to more triggering of fronts of **A** when the active gel was not contracting.

#### 3.15 Figure S15: Influence of the repressor strand $R_A$ concentration on the triggering of a front of A (SI for Figure 4)

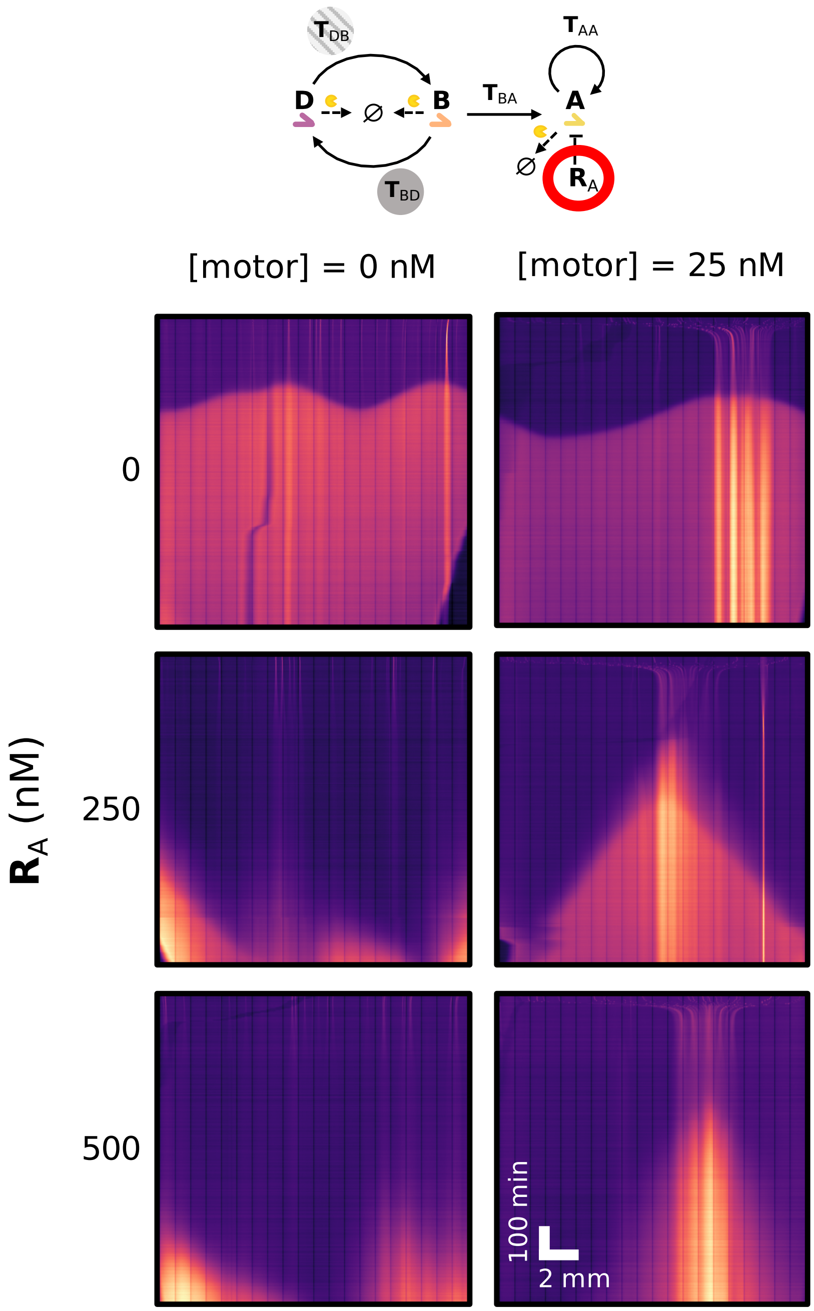

Figure S15: Influence of the repressor strand $\mathbf{R}_A$ concentration on the triggering of a front of **A**. In the absence of the repressor strand $\mathbf{R}_A$, the autocatalytic amplification of **A** was able to start at almost any point of the channel, irrespective of bead activation. We used a repressor at a concentration between 100 and 250 nM to inhibit the self-start of this amplification (*23*). At the highest concentration of repressor tested (500 nM), beads were activated but no front was able to be sustained and to propagate into the channel.

### 4 Supplementary Tables

#### 4.1 Table 1: DNA sequences

Table 1: Sequences of oligonucleotides used in this work. Asterisks * stand for phosphorothioate backbone modifications on the 5' end to protect the strand from being degraded by the exonuclease. All the DNA oligonucleotides were purchased from Biomers.

| Name | Sequence (5'-3') | Modif. 5' | Modif. 3' |
|---|---|---|---|
| A | CGAGTCTGTT | ∅ | ∅ |
| $T_{AA}$ | A*A*CAGACTCGAACAGACTCG | ∅ | Phosphate |
| D | GGAGTCAGTG | ∅ | ∅ |
| G | TGAGTCTTGG | ∅ | ∅ |
| $T_{DB}$ | *C*C*A*AGACTCACACTGACTCC | Biotin-TEG | Phosphate |
| $T_{BD}$ | *C*A*C*TGACTCCCCAAGACTCA | Biotin-TEG | Phosphate or Cy3.5 dye |
| $R_A$ | A*A*A*ACAGACUCG | ∅ | Phosphate |
| $T_{BA}$ | A*A*C*AGACTCGCCAAGACTCA | ∅ | Phosphate |

#### 4.2 Table 2 to Table 5: Composition of the solutions

Table 2: Experimental conditions used for the experiments presented in the main text. Asterisks * indicate that the DNA is put on one side of the channel to trigger the reaction-diffusion front. For the three enzymes involved in the DNA/enzyme subsystem, concentrations are indicated in %(v/v) of the stock solutions. Their stock solution are 10,000 units/mL, 8,000 units/mL and 125 $\mu$M respectively for Bst LF, NBI and ttRecJ.

| Figure | Fig. 2a | Fig. 2b | Fig. 2c | Fig. 3a | Fig. 3b-c, f-g | Fig. 3d-e | Fig. 4c | Fig. 4d | Fig. 4f |
|---|---|---|---|---|---|---|---|---|---|
| Buffer AM (X) | 1 | 1 | 1 | 1 | 1 | 1 | 1 | 1 | 1 |
| Taxol ($\mu$M) | ∅ | ∅ | ∅ | 20 | 20 | 20 | 20 | 20 | 20 |
| Glucose oxidase ($\mu$g/mL) | 150 | 150 | 150 | 150 | 150 | 150 | ∅ | ∅ | ∅ |
| Catalase ($\mu$g/mL) | 25 | 25 | 25 | 25 | 25 | 25 | ∅ | ∅ | ∅ |
| dNTP (mM) | 0.5 | 0.5 | 0.5 | 0.5 | 0.5 | 0.5 | 0.25 | 0.25 | 0.25 |
| Bst LF %(v/v) | 0.1 | **0-0.5** | 0.5 | 0.5 | 0.5 | ∅ | 0.5 | 0.5 | 0.5 |
| NBI %(v/v) | 2.5 | 2.5 | 2.5 | 2.0 | 4.0 | ∅ | 2.5 | 2.5 | 2.5 |
| ttRecJ %(v/v) | ∅ | ∅ | ∅ | ∅ | ∅ | ∅ | 0.125 | **0-0.3** | 0.150 |
| **A** ($\mu$M) | 1* | 1* | 1* | 1* | 1* | ∅ | ∅ | ∅ | ∅ |
| $\mathbf{T}_{AA}$ (nM) | 100 | 100 | 100 | 100 | 100 | ∅ | ∅ | ∅ | 50 |
| **D** (nM) | ∅ | ∅ | ∅ | ∅ | ∅ | ∅ | 5 | 5 | 1 |
| Beads-$\mathbf{T}_{DB}$ (beads/$\mu$L) | ∅ | ∅ | ∅ | ∅ | ∅ | ∅ | 2 | 2 | 0.75 |
| Beads-$\mathbf{T}_{BD}$ (beads/$\mu$L) | ∅ | ∅ | ∅ | ∅ | ∅ | ∅ | 20 | 20 | 5 |
| $\mathbf{R}_A$ (nM) | ∅ | ∅ | ∅ | ∅ | ∅ | ∅ | ∅ | ∅ | 125 |
| $\mathbf{T}_{BA}$ (nM) | ∅ | ∅ | ∅ | ∅ | ∅ | ∅ | ∅ | ∅ | 50 |
| Kinesin (type, nM) | K401, 10 | K401, 10 | K401, **0-20** | K401, 25 | K430, **0-5** | K430, 20 | K430, 15 | K430, **0-15** | K430, **0-25** |
| Microtubules (type, mg/mL) | gmpccp, 0.25 | gmpcpp, 0.25 | gmpcpp, 0.25 | taxol, 0.25 | taxol, 0.5 | taxol, 0.5 | taxol, 0.5 | taxol, 0.5 | taxol, 0.5 |
| Temperature (°C) | 25 | 25 | 25 | 25 | 25 | 25 | 28 | 28 | 28 |

Table 3: Experimental conditions used for the experiments presented in the SI (Figures S1 to S9). Asterisks * indicate that the DNA is put on one side of the channel to trigger the reaction-diffusion front. $^{\times}$dATP, dTTP and dCTP only.

| Figure | Fig. S1 (well-mixed) | Fig. S2 | Fig. S3 | Fig. S4 | Fig. S5 | Fig. S6-S8 | Fig. S9 |
|---|---|---|---|---|---|---|---|
| Buffer AM (X) | **Variable** | 1 | 1 | 1 | 1 | 1 | 1 |
| Taxol ($\mu$M) | ∅ | ∅ | ∅ | ∅ | ∅ | 20 | 20 |
| Glucose oxidase ($\mu$g/mL) | 150 | 150 | 150 | 150 | 150 | 150 | 150 |
| Catalase ($\mu$g/mL) | 25 | 25 | 25 | 25 | 25 | 25 | 25 |
| dNTP$^{\times}$ (mM) | 0.5 | 0.5 | 0.5 | **0-0.5** | 0.5 | 0.5 | 0.5 |
| dGTP (mM) | 0.5 | 0.5 | 0.5 | **0-0.5** | **0-0.25** | 0.5 | 0.5 |
| Bst LF %(v/v) | 0.2 | 0.1 | 0.5 | ∅ | 0.5 | 0.5 | 0.5 |
| NBI %(v/v) | 2.5 | 2.5 | 2.5 | ∅ | 2.0 | 4.0 | 2.0 |
| ttRecJ %(v/v) | ∅ | ∅ | ∅ | ∅ | ∅ | ∅ | ∅ |
| **A** ($\mu$M) | 0.010 | 1* | 1* | ∅ | 1* | 1* | 1* |
| $\mathbf{T}_{AA}$ (nM) | 200 | 100 | 100 | ∅ | 100 | 100 | 100 |
| **D** (nM) | ∅ | ∅ | ∅ | ∅ | ∅ | ∅ | ∅ |
| Beads-$\mathbf{T}_{DB}$ (beads/$\mu$L) | ∅ | ∅ | ∅ | ∅ | ∅ | ∅ | ∅ |
| Beads-$\mathbf{T}_{BD}$ (beads/$\mu$L) | ∅ | ∅ | ∅ | ∅ | ∅ | ∅ | ∅ |
| $\mathbf{R}_{A}$ (nM) | ∅ | ∅ | ∅ | ∅ | ∅ | ∅ | ∅ |
| $\mathbf{T}_{BA}$ (nM) | ∅ | ∅ | ∅ | ∅ | ∅ | ∅ | ∅ |
| Kinesin (type, nM) | ∅ | K401, 10 | K401, **0-20** | K401, 25 | K401, 20 | K430, **0-5** | K401, 5 |
| Microtubules (mg/mL) | ∅ | gmpcpp, 0.25 | gmpcpp, 0.25 | gmpcpp, 0.5 | gmpcpp, 0.5 | taxol, 0.5 | taxol, **0-2** |
| Temperature (°C) | 25 | 25 | 25 | 25 | 25 | 25 | 25 |

Table 4: Experimental conditions used for the experiments presented in the SI (Figures S10 to S15). Asterisks * indicate that the DNA is put on one side of the channel to trigger the reaction-diffusion front. $^{\times}$dATP, dTTP and dCTP only.

| Figure | Fig. S10 | Fig. S11 | Fig. S12-S13 | Fig. S14 | Fig. S15 |
|---|---|---|---|---|---|
| Buffer AM (X) | 1 | 1 | 1 | 1 | 1 |
| Taxol ($\mu$M) | 20 | 20 | 20 | 20 | 20 |
| Glucose oxidase ($\mu$g/mL) | 150 | 150 | ∅ | ∅ | ∅ |
| Catalase ($\mu$g/mL) | 25 | 25 | ∅ | ∅ | ∅ |
| dNTP$^{\times}$ (mM) | 0.5 | 0.5 | 0.25 | 0.25 | 0.25 |
| dGTP (mM) | 0.5 | 0.5 | 0.25 | 0.25 | 0.25 |
| Bst LF %(v/v) | ∅ | ∅ | 0.5 | 0.5 | 0.5 |
| NBI %(v/v) | ∅ | ∅ | 2.5 | 2.5 | 2.5 |
| ttRecJ %(v/v) | ∅ | ∅ | 0.150 | 0.150 | 0.150 |
| **A** ($\mu$M) | ∅ | ∅ | ∅ | ∅ | ∅ |
| $\mathbf{T}_{AA}$ (nM) | ∅ | ∅ | ∅ | 50 | 50 |
| **D** (nM) | ∅ | ∅ | 1 | 1 | 1 |
| Beads-$\mathbf{T}_{DB}$ (beads/$\mu$L) | ∅ | ∅ | 2 | **0-2** | 0.75 |
| Beads-$\mathbf{T}_{BD}$ (beads/$\mu$L) | ∅ | ∅ | 20 | 5 | 5 |
| $\mathbf{R}_{A}$ (nM) | ∅ | ∅ | ∅ | 125 | **0-500** |
| $\mathbf{T}_{BA}$ (nM) | ∅ | ∅ | ∅ | 50 | 50 |
| Kinesin (type, nM) | K430, 20 | K430, 20 | K430, 20 | K430, 25 | K430, 25 |
| Microtubules (mg/mL) | taxol, 0.5 | taxol, 0.5 | taxol, 0.5 | taxol, 0.5 | taxol, 0.5 |
| Temperature (°C) | 25 | 25 | 28 | 28 | 28 |

Table 5: Experimental conditions used for the experiments presented in the movies. Asterisks * indicate that the DNA is put on one side of the channel to trigger the reaction-diffusion front. $^{\times}$dATP, dTTP and dCTP only.

| Figure | Movie S1 | Movie S2 | Movie S3 | Movie S4 | Movie S5 | Movie S6 | Movie S7 |
|---|---|---|---|---|---|---|---|
| Buffer AM (X) | 1 | 1 | 1 | 1 | 1 | 1 | 1 |
| Taxol ($\mu$M) | ∅ | ∅ | 20 | 20 | 20 | 20 | 20 |
| Glucose oxidase ($\mu$g/mL) | 150 | 150 | 150 | 150 | 150 | ∅ | ∅ |
| Catalase ($\mu$g/mL) | 25 | 25 | 25 | 25 | 25 | ∅ | ∅ |
| dNTP$^{\times}$ (mM) | 0.5 | 0.5 | 0.5 | 0.5 | 0.5 | 0.25 | 0.25 |
| dGTP (mM) | 0.5 | 0.05 | 0.5 | 0.5 | 0.5 | 0.25 | 0.25 |
| Bst LF %(v/v) | 0.1 | 0.5 | 0.5 | 0.5 | ∅ | 0.5 | 0.5 |
| NBI %(v/v) | 2.5 | 2.0 | 2.0 | 2 | ∅ | 2.5 | 2.5 |
| ttRecJ %(v/v) | ∅ | ∅ | ∅ | ∅ | ∅ | 0.125 | 0.150 |
| **A** ($\mu$M) | 1* | 1* | 1* | 1* | ∅ | ∅ | ∅ |
| $\mathbf{T}_{AA}$ (nM) | 100 | 100 | 100 | 100 | ∅ | ∅ | 50 |
| **D** (nM) | ∅ | ∅ | ∅ | ∅ | ∅ | 5 | 1 |
| Beads-$\mathbf{T}_{DB}$ (beads/$\mu$L) | ∅ | ∅ | ∅ | ∅ | ∅ | 2 | 0.75 |
| Beads-$\mathbf{T}_{BD}$ (beads/$\mu$L) | ∅ | ∅ | ∅ | ∅ | ∅ | 20 | 5 |
| $\mathbf{R}_{A}$ (nM) | ∅ | ∅ | ∅ | ∅ | ∅ | ∅ | 125 |
| $\mathbf{T}_{BA}$ (nM) | ∅ | ∅ | ∅ | ∅ | ∅ | ∅ | 50 |
| Kinesin (type, nM) | K401, 10 | K401, 20 | K401, 1 | K401, 5 | K430, 20 | K430, 0-15 | K430, 0-25 |
| Microtubules (mg/mL) | gmpccp, 0.25 | gmpcpp, 0.50 | taxol, 0.5 | taxol, 0.25 | taxol, 0.5 | taxol, 0.5 | taxol, 0.5 |
| Temperature (°C) | 25 | 25 | 25 | 25 | 25 | 28 | 28 |

### 5 Legends of Movies S1 to S7

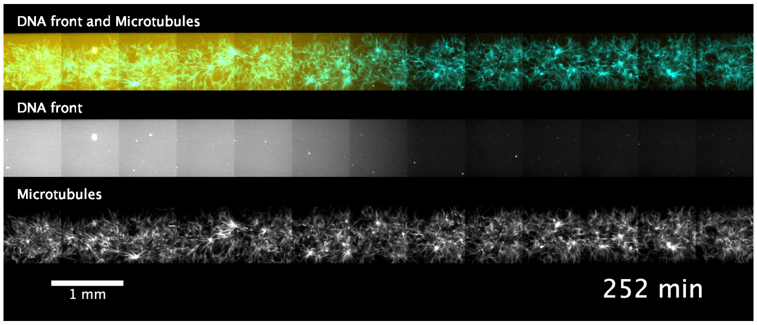

Movie S1: Propagation of the DNA front in an active gel that displays a locally contractile behavior. Associated to Figure 1 (d).

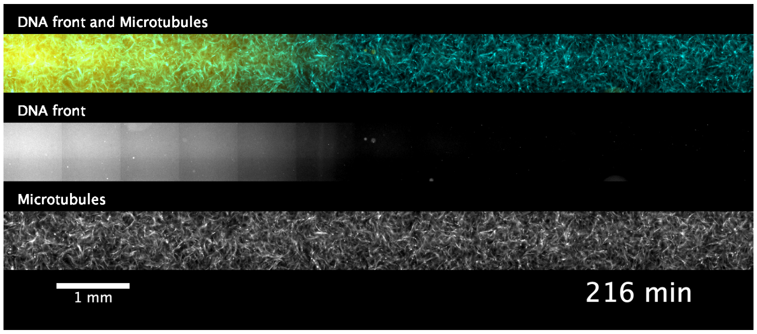

Movie S2: Propagation of the DNA front in an active gel that displays chaotic flows. In the presence of GMPCPP-microtubules and a relatively low concentration of dGTP ($< 50$ $\mu$M), the active fluid displays chaotic flows that slow down after 300 min to eventually form microtubules aggregates.

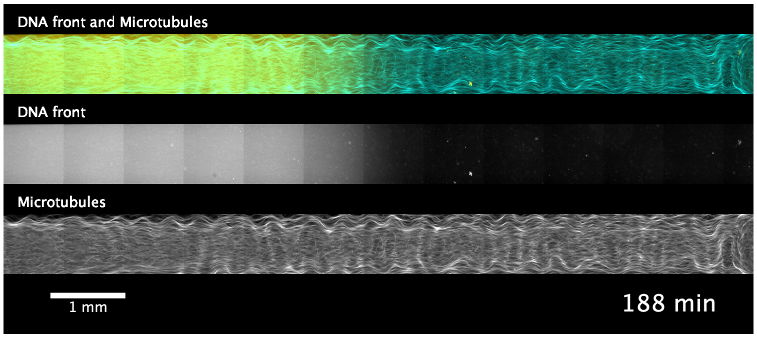

Movie S3: Propagation of the DNA front in an active gel that displays corrugated patterns. In the presence of longer, taxol-stabilized microtubules and a relatively low concentration of kinesin motors ($< 2.5$ nM), the active gel formed a corrugated sheet that undulated in the $xz$ plane.

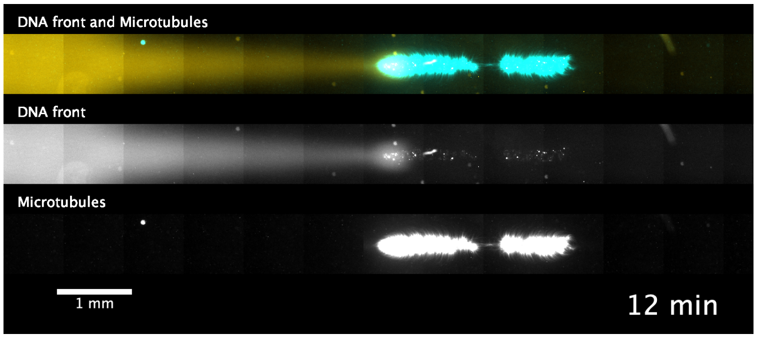

Movie S4: A globally contracting active gel perturbs and accelerates the propagation of the DNA front. Associated to Figure 3 (a).

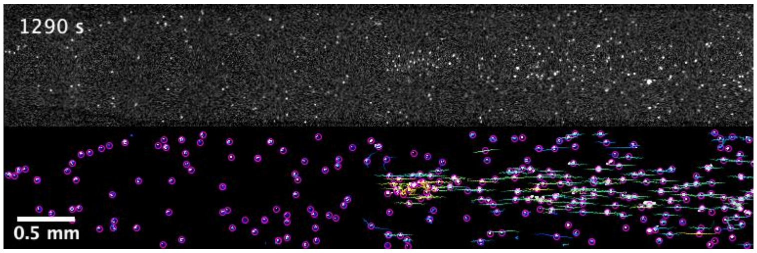

Movie S5: Visualization of the hydrodynamic flow induced by the global contraction of the active gel by using passive fluorescent beads. (Top) Raw fluorescent images. (Bottom) Tracking of the fluorescent beads. The local velocity direction is displayed as small segments. The gel corresponds to the region where the beads move from left to right. Associated to Figure 3 (d).

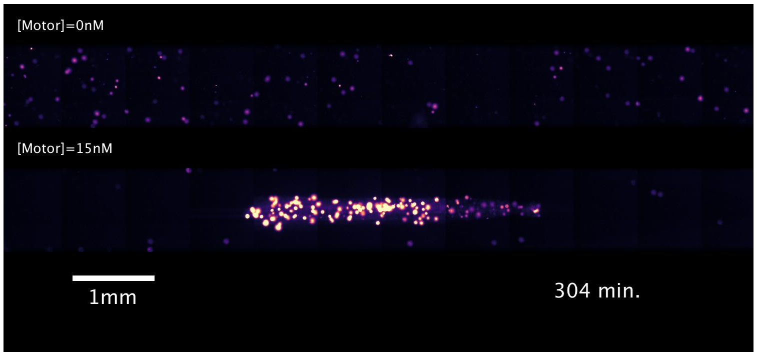

Movie S6: A globally contracting active gel triggers the activation of a cross-autocatalytic reaction localized on hydrogel beads. Associated to Figure 4 (c).

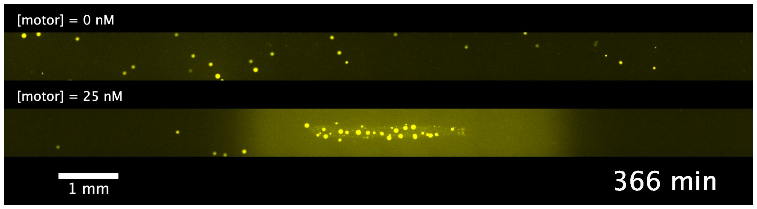

Movie S7: A globally contracting active gel triggers the activation of a cross-autocatalytic reaction localized on hydrogel beads and a DNA front in solution. Associated to Figure 4 (f).
